# The ClpX protease is essential for removing the CI master repressor and completing prophage induction in *Staphylococcus aureus*

**DOI:** 10.1101/2022.09.18.507959

**Authors:** Mohammed A. Thabet, José R. Penadés, Andreas F. Haag

**Affiliations:** School of Infection & Immunity, University of Glasgow, G12 8TA, Glasgow, UK; Department of Biology, Faculty of Science and Art in Al-Mandq, Al-Baha University, Al-Baha city, 576J+R9 Musaylah, Kingdom of Saudi Arabia; MRC Centre for Molecular Bacteriology and Infection, Imperial College London, SW7 2AZ, UK; School of Medicine, University of St Andrews, North Haugh, St Andrews, KY16 9TF, UK

**Keywords:** Prophage induction, ClpX, ClpP, SOS response, phage cycle

## Abstract

Bacteriophages (phages) are the predominant biological entities on the planet and play an important role in the spread of bacterial virulence, pathogenicity, and antimicrobial resistance. After infection, temperate phages can integrate in the bacterial chromosome thanks to the expression of the prophage-encoded CI master repressor. Upon SOS induction, and promoted by RecA*, CI auto-cleaves generating two fragments, one containing the N-terminal domain (NTD), which retains strong DNA-binding capacity, and other corresponding to the C-terminal part of the protein. However, it is unknown how the CI NTD is removed, a process that is essential to allow prophage induction. Here we identify for the first time that the specific interaction of the ClpX protease with the CI NTD repressor fragment is essential and sufficient for prophage activation after SOS-mediated CI autocleavage, defining the final stage in the prophage induction cascade. Our results provide unexpected roles for the bacterial protease ClpX in phage biology.

## Introduction

Bacteriophages, or phages, are the most abundant biological entity on the planet (1) and crucial agents of horizontal gene transfer, including the dissemination of antimicrobial resistance genes and virulence factors (2). Temperate phages, such as the model phage λ, have established an intricate relationship with their bacterial host where they can either replicated and kill their host cell (lytic cycle) or integrate into their host’s genome and be passed on vertically (lysogenic cycle) until they become reactivated upon defined environmental cues. Lysogeny is controlled by a phage repressor, called CI in λ, which represses the transcription of genes required for phage excision, replication, packaging, and host cell lysis as well as its own transcription (3,4). Upon induction, the prophage produces the lytic regulator protein Cro, which is encoded divergent to the *c*I repressor gene, binds to the *c*I promoter, and stabilizes the lytic cycle by blocking transcription from the P_RM_ promoter driving *c*I transcription (3). Expression levels of CI and Cro are finely balanced to ensure tight repression of the phage lytic cycle while at the same time enabling its rapid activation upon sensing the correct stimulus.

The bacterial SOS response is one of the key signals sensed to activate the phage lytic cycle. The SOS response pathway is triggered by DNA damage and coordinates the expression of genes involved in DNA repair, replication, and cell division (5,6). In the absence of DNA damage, the LexA repressor binds to operator sequences within the promoter of SOS-response genes (SOS boxes/Cheo boxes in *Bacillus subtilis*) (7,8). LexA binds to these boxes as a dimer with each monomer comprising a C-terminal dimerization domain (CTD) and an N-terminal DNA-binding domain (NTD) (9). DNA damage results in the generation of single stranded DNA fragments to which RecA binds resulting in an activated RecA-nucleoprotein filaments (RecA*) (10). RecA* in turn triggers the autocatalytic cleavage of unbound LexA (11), reducing the cellular LexA pool and allowing the progressive activation of SOS-genes according to their LexA binding affinities. Cleavage of the LexA repressor separates its CTD and NTD, revealing latent ClpX-recognition motifs leading to the degradation of both fragments by the ClpXP protease complex (12). Even though both fragments of the *E. coli* LexA protein are recognized by ClpX, ClpX is mainly involved in the degradation of the LexA NTD *in vivo* (12), while the Lon protease is responsible for degrading the LexA CTD (13). In *Staphylococcus aureus*, the ATPases ClpX and ClpC as well as the proteolytic activity of ClpP have been implicated in the degradation of the LexA NTD and subsequent SOS activation (14). Since the LexA NTD retains some of its ability to bind to SOS boxes and to repress target genes in both Gram-positive and Gram-negative bacteria (14–16), its targeted removal by ClpXP is important for SOS activation.

Prophage induction is a fundamental process in phage biology and is controlled by the CI protein in phage λ. λ CI is a LexA-like repressor comprised of an NTD involved in DNA binding and repression and a catalytic C-terminal dimerization domain that upon recognition of activated RecA* are autocatalytically separated (17). CI repressors are present in a wide variety of temperate phage genomes of both Gram-positive and Gram-negative hosts (18,19). Interestingly, several examples of phage CI repressor molecules have been identified that retain some and sometimes even full binding affinities with their cognate promoter DNA after autocatalytic separation of their C- and NTDs (20–23). The retention of such levels of DNA affinity by the CI NTD therefore would require its active elimination to ensure progression from the lysogenic to the lytic phage lifecycle. Despite its vital role in temperate phage biology, how the CI NTD is eliminated after RecA* mediated release remains elusive.

The proteolytic degradation of the LexA NTD together with the conserved domain architecture found in both LexA and LexA-like repressors such as CI suggest that the bacterial proteasome and specifically Clp proteases might also be involved in prophage induction. These proteases are key mediators of the ATP-dependent degradation of proteins and are usually constituted of a proteolytic subunit (ClpP or ClpQ/HslV) which associates with an AAA-ATPase (ClpX, ClpA, ClpC and ClpE with ClpP; ClpY/HslU with ClpQ/HslV) (24). ClpP alone is only able to degrade small peptides and requires association with one of its ATPases which act as energy-dependent unfoldases, unfolding and delivering the protein into the ClpP proteolytic chamber (25). Target specificity of the protease complex is mediated by its ATPase, which can recognize specific degron motifs either alone or in association with additional adaptor proteins (24,26–28).

In this study we identify that the ClpX ATPase performs an essential role in prophage induction by removing the CI NTD after RecA* mediated CI cleavage and allowing the prophage to enter lytic replication. This process is distinct from the role of ClpX on the staphylococcal SOS response and acts via direct binding of ClpX to the CI NTD. By contrast, the ClpP protease and its proteolytic interaction with ClpX are not required for elimination of the CI NTD but are essential for the activation of the staphylococcal SOS response.

## Materials and Methods

### Bacterial strains and culture conditions

The bacterial strains used in this study are listed in Table S2. *S. aureus* strains were grown in Tryptic soy broth (TSB) or on Tryptic soy agar (TSA) plates. *E. coli* strains were grown in Luria–Bertani broth (LB) and on LB agar plates. Antibiotic selection was used where appropriate (10 or 2.5 μg ml^−1^ erythromycin (as indicated), 100 μg ml^−1^ ampicillin and 30 μg ml^−1^ kanamycin).

### Phage infection assay

Test strains were grown in TSB to early exponential phase (OD_540_~0.15) at 37°C and 120 rpm. 12 ml of this culture were pelleted by centrifugation (3300 × g, 10 min) and the pellet was resuspended in 6 ml of fresh TSB and 6 ml of phage buffer (1 mM MgSO_4_, 4 mM CaCl_2_, 50 mM Tris-Cl, 100 mM NaCl, pH=8). Equal numbers of phages (~10^8^ PFU) were added to each culture and infected cultures incubated for 4 h at 30°C and 80 rpm followed by overnight incubation at room temperature. Cleared lysates were filtered using a 0.2 μm syringe filter (Satorius).

### Phage induction and titration

*S. aureus* strains lysogenic for the required phages were grown to early exponential phase (OD_540_~0.15) at 37°C and 120 rpm. Cultures were then induced by the addition of mitomycin C (2 μg ml^−1^) and incubated for 4-5 h at 30°C, 80 rpm followed by overnight incubation at room temperature before filtering with a 0.2 μm syringe filter (Sartorius). To determine the phage titers, RN4220 cultures were grown to OD_540_~0.35 and 100 μl of this culture were mixed with 100 μl of serially diluted phage lysates in phage buffer (1 mM MgSO_4_, 4 mM CaCl_2_, 50 mM Tris-Cl, 100 mM NaCl, pH=8). The mixtures were incubated for 5 min at room temperature, then 3 ml of phage top agar (PTA, 20 g l^−1^ Nutrient Broth No. 2, Oxoid, plus 3.5 g l^−1^ agar, Formedium supplemented with 10 mM CaCl_2_) was added and the mixture overlaid onto phage base agar plates (20 g l^−1^ Nutrient Broth No. 2, Oxoid, plus 7 g l^−1^ agar, Formedium supplemented with 10 mM CaCl_2_). Plates were incubated overnight at 37°C and plaque forming unit (PFU ml^−1^) determined.

### Superinfection exclusion / Efficiency of plating assay

*S. aureus* strains carrying the required plasmids were grown in TSB supplemented with the relevant antibiotics to an OD_540_~0.35 and 100 μl of this culture were mixed with 3 ml of phage top agar (PTA, 20 g l^−1^ Nutrient Broth No. 2, Oxoid, plus 3.5 g l^−1^ agar, Formedium supplemented with 10 mM CaCl_2_ and 1 μM CdCl_2_) and overlaid onto phage base agar plates (20 g l^−1^ Nutrient Broth No. 2, Oxoid, plus 7 g l^−1^ agar, Formedium supplemented with 10 mM CaCl_2_ and 1 μM CdCl_2_). Phage lysates and dilutions in phage buffer (1 mM MgSO_4_, 4 mM CaCl_2_, 50 mM Tris-Cl, 100 mM NaCl, pH=8) were spotted in triplicates of 10 μl each onto lawns of the specified strains, dried and incubated overnight at 37°C prior to plaque forming unit (PFU ml^−1^) determination.

### DNA manipulations

General DNA manipulations were performed using standard procedures. Plasmid constructs used in this study (Table S3) were generated by cloning PCR products (Kapa Hifi Polymerase, Roche) obtained with oligonucleotide primers listed in Table S4 and digested with the indicated restriction enzymes (New England Biolabs). Detection probes for phage DNA in Southern blots were generated by PCR using a non-proofreading polymerase (DreamTaq polymerase, ThermoFisher) using oligonucleotides specified in Table S4. Probe labelling and DNA hybridization were performed following the protocol provided with the PCR-DIG DNA-labelling and chemiluminescent detection kit (Roche).

### Generation of clean deletion mutants

The required clean deletion mutants were constructed using allelic replacement by cloning flanking regions up- and downstream of the respective gene (0.5-1 kbp) into pMAD (29) using oligonucleotide primers described in Table S4. The plasmids were then transformed into the required strains and integration of the plasmid was selected by growth at the restrictive temperature (42°C) on TSA plates supplemented with 80 μg ml^−1^ X-gal and 2.5 μg ml^−1^ erythromycin. Single crossover events were isolated (light blue colonies) and grown overnight under replication-permissive conditions (TSB, 30°C, 80 rpm) to facilitated excision and loss of the integrated plasmid. Serial dilutions of the cultures were plated on TSA plates supplemented with 80 μg ml^−1^ X-gal and correct deletion mutants identified by PCR followed by sequencing using oligonucleotides annealing outside of the recombination region and indicated in Table S4.

### Southern blotting

Strains containing the defined phage and plasmids were grown to early exponential phase (OD_540_~0.15) in 10 ml of TSB supplemented with antibiotics where plasmids were present. Phages were induced with mitomycin C (2 μg ml^−1^) and, where pCN51 expression plasmid derivatives were present, 1 μM CdCl_2_ was added to induce expression as indicated. One ml samples were taken at the defined timepoints, pelleted by centrifugation (16873 × g, 2 min) and pellets shock frozen on dry ice. The pellets were re-suspended in 50 μl lysis buffer (47.5 μl TES-Sucrose (10 mM Tris-Cl, 100 mM NaCl, 1 mM EDTA, 20% (w/v) sucrose) and 2.5 μl lysostaphin [12.5 μg ml^−1^]) and incubated at 37°C for 1 h. Following the incubation, 55 μl of SDS 2% proteinase K buffer (47.25 μl H_2_O, 5.25 μl SDS 20%, 2.5 μl proteinase K [20 mg ml^−1^]) was added before incubation at 55°C for 30 min. Samples were vortexed for 1 h with 11 μl of 10x loading dye followed by three cycles of 5 min incubations in dry ice/ethanol and at 65°C in a water bath. Samples were run on 0.7% agarose gel at 25-30 V overnight. DNA was transferred by capillary action to a positively charged nylon membrane (Roche), processed as per the manufacturer’s instructions, and exposed using a DIG-labelled probe (see DNA methods) and anti-DIG antibody (1:10000 (v/v), Roche, product 11093274910) before washing and visualization.

### Two-hybrid assay

The two-hybrid assay for protein-protein interaction was performed as described previously (30,31) using two compatible plasmids: pUT18c expressing T18 fusions with either ClpX_WT_ or ClpX_I265E_ and pKT25 expressing the T25 fusion with either the Φ11 CI_WT_ or its NTD CI_G131*_. Both plasmids were co-transformed into *E. coli* BTH101 for the Bacterial Adenylate Cyclase Two Hybrid (BACTH) system and plated on LB supplemented with ampicillin (100 μg ml^−1^), kanamycin (30 μg ml^−1^), X-gal (20 μg ml^−1^) and IPTG (100 μM). After incubation at 30°C for 48 h (early reaction) to 72 h (late reaction), the protein-protein interaction was detected by a color change. Blue colonies represent an interaction between the two clones, while white/yellow colonies are negative for any interaction.

### Promoter activity assay

For the β-lactamase assays, overnight cultures were prepared by inoculating a single colony from each strain into a 5 ml TSB supplemented with 10 μg ml^−1^ erythromycin and incubated at 37°C, 120 rpm for 16-18 h. The cultures were then diluted 1/50 in 13 ml of fresh TSB supplemented with 10 μg ml^−1^ erythromycin and grown at 37°C and 120 rpm to early exponential phase (OD_540_~0.15-0.2). 200 μl of culture were added directly to 800 μl of potassium phosphate buffer (50 mM, pH 5.9, supplemented with 10 mM sodium azide) and frozen on dry ice. Where SOS-induction of the *cro* promoter was assessed, cultures were split in two (6 ml each), induced either with or without MC (2 μg ml^−1^) and incubated at 30°C, 80 rpm until sampling as described above. β-lactamase assays, using nitrocefin as substrate, were performed as described (32,33): 50 μl of the collected sample were mixed with 50 μl of nitrocefin stock solution (192 μM made in 50 mM potassium phosphate buffer, pH 5.9), and immediately reading the absorbance at 490 nm using an ELx808 microplate reader (BioTek) for 30 min. Promoter activity was calculated as Promoter activity = (*d*A_490_/*d*t(h))/(OD_540_ × d × V), where OD_540_ is the absorbance of the sample at OD_540_ at collection, d is the dilution factor, and V is the sample volume. Note that only the linear segment of the resulting absorbance readings is considered for activity calculations.

### Preparation of samples and quantitative whole genome sequencing

Overnight cultures were prepared by inoculating a single colony from each strain into a 5 ml TSB and incubated at 37°C, 120 rpm for 16-18 h. The desired strains were then diluted 1:50 in 13 ml of fresh TSB and grown to exponential phase (OD_540_~0.15-0.20) to collect samples before induction. Next, the cultures were treated with MC (2 μg ml^−1^) and incubated at 30°C and 80 rpm for 60 min prior to sample collection. One ml samples were collected, and genomic DNA was extracted using the GenElute Bacterial DNA kit (Sigma Aldrich) according to the manufacturer’s instructions. The DNA was precipitated by adding 10% (v/v) 3 M sodium acetate (pH 5.2), 2.5 volumes of 100% ethanol and incubation of the mixture for 1 h at −80°C. The DNA was then pelleted at 16873 × g for 30 min at 4 °C and washed once with 1 ml of ice cold 70% (v/v) ethanol. After centrifugation, the DNA pellets were air dried for 30 min and resuspended in 25 μl of TE buffer (10 mM Tris-HCl, 1 mM EDTA, pH 8.0). Quality control of DNA samples was tested using Agilent Bioanalyzer 2100 and whole genome sequencing (WGS) was performed at the University of Glasgow’s Polyomics Facility using Illumina TruSeq DNA Nano library prep, obtaining 2 × 75 bp pair end reads with DNA PCR free libraries.

### Preparation of custom reference genomes

Genomic DNA of the RN450 (NCTC-4) reference stock in the lab (JP1250) was extracted and sequenced as described above. Next, reads were assembled to a scaffold of the deposited NCTC8325 (GenBank Accession CP000253) reference genome using the PATRIC Bioinformatics Resource Center (34). The three prophages of NCTC8325 were deleted and any mutations identified were curated manually. Sequencing reads were then reassembled to the curated genome (GenBank Accession CP097113) as described above for verification. The sequences of either Φ11 (RefSeq Accession NC_004615) or 80α (RefSeq Accession NC_009526) were inserted into attachment sites 5 and 7 (35), respectively and correct insertion verified by assembly of genome sequencing reads for strain JP18269 (GenBank Accession CP097114) or JP18270 (GenBank Accession CP097115) for Φ11 or 80α lysogens, respectively. The curated genomes were next uploaded to the Galaxy web platform, and we used the public server at usegalaxy.org to analyze the data (36). Genomes were reannotated using Prokka v1.14.6 (37,38) (Galaxy Version 1.14.6+galaxy1).

### Analysis of whole genome sequencing data

The sequencing data were uploaded to the Galaxy web platform, and we used the public server at usegalaxy.org to analyze the data (36). The read quality of paired reads was assessed using FastQC v0.11.8 (39) (Galaxy Version 0.72+galaxy1) followed by adapter trimming using Trimmomatic v0.38 (40) (Galaxy Version 0.38.0) and standard setting for paired end reads and Illumina data. Trimmed reads were then reassessed using FastQC and mapped to custom genomes of RN450 reference genome containing either prophage Φ11 or 80α using the Burrows-Wheeler Alignment Tool v0.7.17.4 (41,42) (Galaxy Version 0.7.17.4) with default settings and saved as BAM files. To normalize sequence coverage across experiments, we first filtered the aligned reads mapping to the bacterial chromosome and not belonging to the prophage using the BAMTools v2.4.0 Filter tool (43) (Galaxy Version 2.4.1). The number of mapped reads for each experiment were extracted from the filtered BAM files using the SAMTools stats utility v1.9 (44) (Galaxy Version 2.0.2+galaxy2). Average genome coverage was calculated using the following formula: average genome coverage = (number of mapped reads) × (average read length (bp)) / (genome length (bp)). Next, we computed the relative coverage over 50 bp sliding windows along the entire chromosome without normalization for each of the experiments of the unfiltered BAM files using the bamCoverage tool of the deepTools2 package v3.3.2 (45) (Galaxy Version 3.3.2.0.0). These coverage files were saved in bedgraph format and further analyzed using RStudio v2021.9.1.372 (Ghost Orchid) (46) and R v4.1.2 (47). Samples were normalized by dividing each coverage window by the average genome coverage calculated for each experiment. Final coverage graphs were plotted in RStudio using ggplot2 v3.3.5 (48) and genome organization around the plotted area extracted from the gff3 file produced by the Prokka annotation and graphed using gggplo2 and the gggenes package v0.4.1 (49).

### Statistical analyses

Statistical analysis was performed as indicated in the figure legend. In general, phage titers were log_10_-transformed and analyzed by either One-Way ANOVA followed by Tukey’s HSD post-test or using a Student’s unpaired two-tailed t-test as appropriate for the relevant comparison. Promoter activity data were analyzed on raw activity data by either One-Way ANOVA followed by Tukey’s HSD post-test or using a Student’s unpaired two-tailed t-test as appropriate for the relevant comparison. All analysis was done using RStudio.

## Results

### Clp proteases are not involved in the lytic phage cycle

We initiated this study testing the possibility that the chromosomally encoded Clp ATPases or proteolytic subunits (Table S1) could control the life cycles of the prototypical *S. aureus* phages Φ11 and 80α, which have been extensively used as models for Gram-positive phages (50,51). Both phages encode a CI-like repressor that maintains the phage in its lysogenic prophage state within the bacterial chromosome. To test whether any of the Clp proteases (B, C, L, P, Q, X or Y; **Error! Reference source not found**.) were involved in the reproduction of phages Φ11 and 80α, we initially evaluated the lytic cycle of these phages. Lysates of either phage Φ11 and 80α were used to infect strain RN450 or a collection of derivative mutant strains in which the different *clp* genes were either deleted or inactivated by the insertion of an erythromycin resistance cassette. No differences in either the number or in the size of the phage plaques compared to the wt staphylococcal strain were observed when the different mutant strains were infected (Fig. S1). Since plaque formation requires normal phage replication, packaging and lysis, this result suggests that the Clp proteases were not required in these steps of the phage cycles.

### ClpP and ClpX are involved in prophage induction in *S. aureus*

However, there is an additional stage in the cycle of the temperate phages, induction, that could be controlled by any of these proteases. To address this point, we initially used the aforementioned collection of RN450 (non-lysogenic) derivatives, in which the different *clp* genes were either deleted or disrupted. These mutant strains were lysogenized with phages Φ11 or 80α and the prophages were SOS induced by addition of mitomycin C (MC). Next, the number of phages present in the different lysates were analyzed using strain RN4220 as recipient. All strains apart from the *clp*X and *clp*P mutants lysed 4 h after induction of the SOS response, suggesting ClpX and ClpP were involved in prophage induction. Indeed, the phage titer obtained after induction of the prophage present in the *clp*P mutant was reduced 10^5^- to 10^6^-fold (Φ11 and 80α, respectively) compared to that observed for when the prophage was induced from the wt strain, while no plaques were obtained from the prophage present in the *clp*X mutant strain (Fig. S1), indicating that while ClpP was extremely important for prophage induction, ClpX was essential. Note that even in strains that are insensitive to the SOS response, either by carrying mutations in *rec*A or by expressing an insensitive LexA protein, there is always some basal induction of the resident prophages (52–55), an effect that is not seen in the *clp*X mutant.

To clearly confirm the previous results, we generated a new set of in-frame deletion mutants in which the *clp*X and *clp*P genes were deleted using allelic replacement (strains JP18031 and JP18030, respectively). These strains were either infected with Φ11 or 80α or lysogenized with these phages (JP18157: Φ11 Δ*clp*X; JP18158: Φ11 Δ*clp*P, JP18169: 80α Δ*clp*X; JP18170: 80α Δ*clp*P) and the phage cycle MC-induced. In agreement with the previous results, phage titers were unaffected during phage infection (Fig. 1A) but showed significant reductions in phage titers after prophage induction (Fig. 1B). As with the insertional mutants, the clean deletion mutants did not lyse after 4 h and only the Δ*clp*P mutant lysed after overnight incubation while the Δ*clp*X mutant lysates remained turbid (Fig. S2). Consistent with this, phages were virtually absent in the Δ*clp*X and reduced ~10^4^-fold in the Δ*clp*P mutant. Introducing plasmid-borne versions of either *clp*X (pJP2601, JP18381 or JP18919) or *clp*P (pJP2602, JP18383 or JP18921) into the respective mutants under the control of a cadmium-inducible promoter fully restored wild type phage lysis and titers (Fig. 1C) confirming their role in prophage induction.

**Fig. 1.**
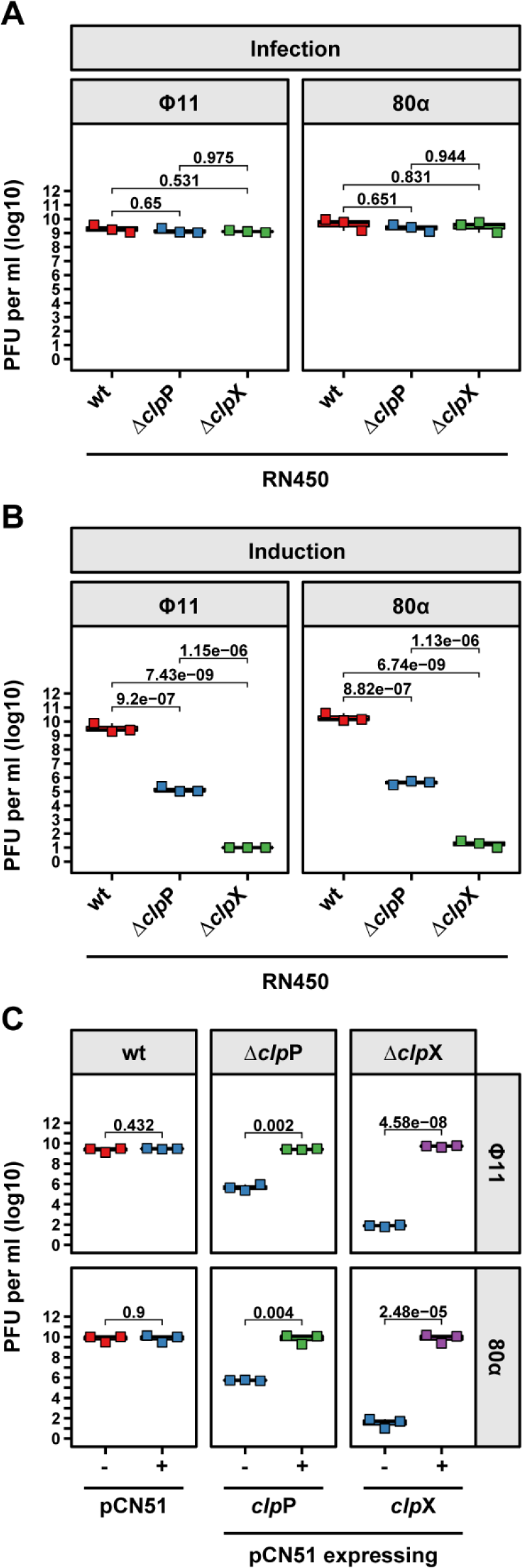
ClpP and ClpX are involved in phage induction but not phage infection. The defined RN450 derivative strains were either **(A)** infected with the indicated phages or **(B)** lysogenic derivatives thereof induced by MC addition. **(C)** The *clp*P and *clp*X genes were cloned into the cadmium-inducible expression plasmid pCN51, introduced into the defined strains, and induced by MC addition. Expression from the pCN51 plasmids was maintained throughout the experiment by the addition of 1 μM CdCl_2_. **(A-C)** Plaque formation was assessed on a lawn of RN4220. Bold horizontal lines in each boxplot represent the median and lower and upper hinges the first and third quartiles, respectively (n=3 biological replicates). Assessment of statistically significant differences between groups was performed using ANOVA followed by Tukey’s HSD post-test **(A&B)** or a two-sided Student’s t-test **(C)** on log_10_ transformed data. p-values are indicated above the respective comparison.

### ClpX and ClpP have different roles in SOS and prophage induction

Since phage induction requires the bacterial SOS response, and since in *S. aureus* this process is affected in the absence of the ClpXP proteases (14), we tested whether the impact of the *clp*X and *clp*P mutations on phage induction could be explained by their role on SOS response activation. This was unlikely, however, since both mutants showed different behaviors and because of the unique phenotype observed with the *clp*X mutant. For this we constructed a set of reporter plasmids in which the promoters of the SOS-inducible genes *lex*A and *rec*A were fused to a β-lactamase reporter gene in plasmid pCN41 (pJP2596 and pJP2597, respectively). These were introduced into the RN4220 wt, RN4220 Δ*clp*P and RN4220 Δ*clp*X mutant strains and the SOS response induced by MC addition (Fig. 2). Consistent with their SOS-inducibility, both *lex*A and *rec*A promoters were activated upon MC addition in the wt *S. aureus* strain background (JP20858 and JP20417, respectively) showing similar fold induction changes (Fig 2A&B, respectively). No activity of either the *lex*A or *rec*A promoters, neither with nor without MC addition, was observed in the Δ*clp*P mutant strain (JP20860 and JP20419, respectively), showing that ClpP was essential for the activation of the SOS response. This confirmed that ClpP-mediated degradation of the LexA NTD was required for SOS-induction. To confirm the essentiality of ClpP to activate the SOS response, we introduced the *lex*A and *rec*A reporter plasmids into an RN4220 derivative strain expressing an SOS-insensitive version of the LexA repressor (LexA_G94E_) that can no longer catalyse auto-cleavage and consequently can no longer induce the SOS response (56,57) (JP20859 and JP20418, respectively). As expected, neither reporter was inducible by MC addition and expression levels were similar to those observed in the Δ*clp*P mutant background.

**Fig. 2.**
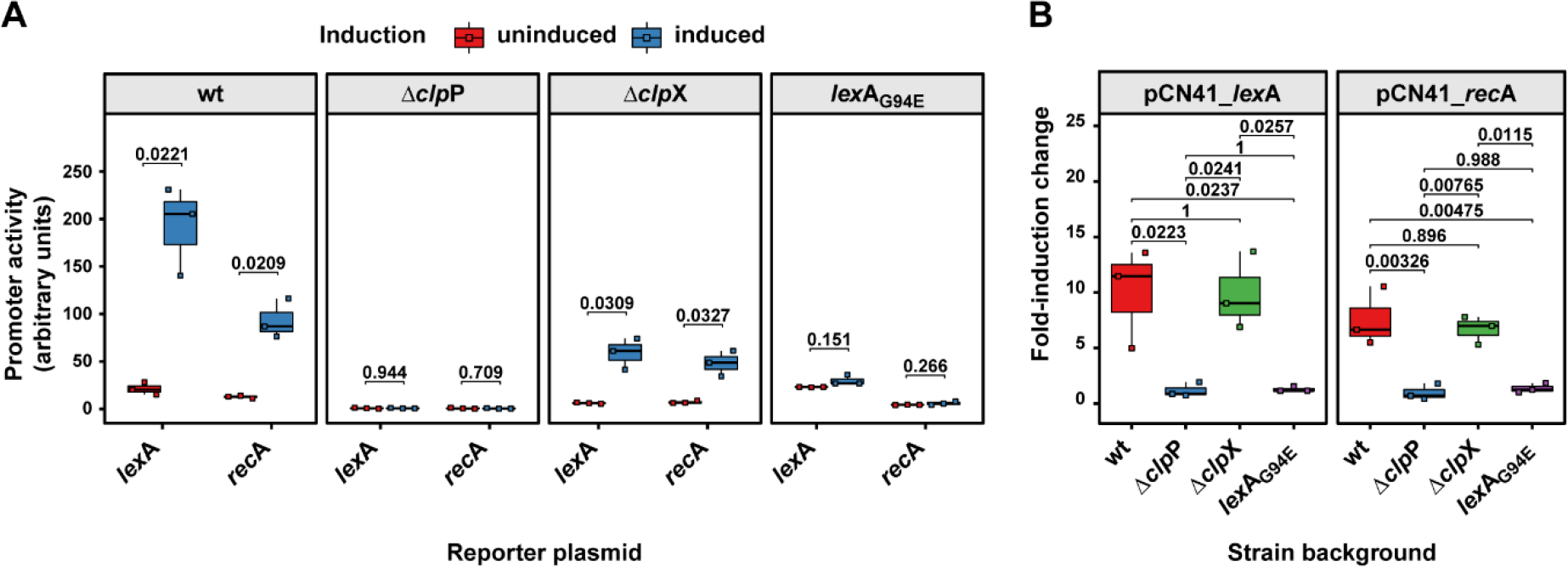
Distinct roles for ClpP and ClpX in SOS response induction. Reporter plasmids were designed to place the β-lactamase reporter gene (*bla*Z) of plasmid pCN41 under the control of the SOS-controlled promoters of *lex*A or *rec*A. **(A)** RN4220 derivative strains containing the indicated plasmids were grown to exponential phase, split and the SOS response induced in one half of the culture with MC (blue boxplots) while the other half was left untreated (red boxplots). Samples were taken 90 min after induction. **(B)** Fold-induction change of MC-induced against non-induced samples. Bold horizontal lines in each boxplot represent the median and lower and upper hinges the first and third quartiles, respectively (n=3 biological replicates). Assessment of statistically significant differences between groups was performed using **(A)** a two-sided Student’s t-test or **(B)** ANOVA followed by Tukey’s HSD post-test. p-values are indicated above each comparison.

By contrast, when the *lex*A and *rec*A reporter plasmids were introduced in the Δ*clp*X mutant background (JP20861 and JP20420, respectively), although the transcriptional levels of both *lex*A and *rec*A reporters were reduced, they could still be induced in the presence of MC with a comparable fold-induction change to the wt strain (Fig 2A&B, respectively), suggesting that ClpX was involved but was not essential in inducing the staphylococcal SOS response.

Taken together, the impact of the *clp*P and *clp*X mutation in prophage induction and on the activation of the SOS response via *lex*A and *rec*A expression suggest that their role in both processes is different.

### ClpX is essential for phage replication

The previous results clearly showed that ClpP and ClpX were involved in both prophage and SOS response induction. However, while ClpX was essential for prophage but not for SOS-response induction, ClpP was vital for the induction of the staphylococcal SOS-response and highly important for prophage activation. The different roles of ClpP and ClpX in SOS- and prophage-induction prompted us to further explore their function in prophage induction. Reduced prophage titres can be the result of a failure in phage induction, replication, packaging or host cell lysis and progeny release. Since our previously results demonstrated that these proteins were not required during the lytic cycle of the phage, we reasoned that the deficiencies observed in the Δ*clp*P and Δ*clp*X mutants should be related with the inability of these mutants to eliminate the CI-mediated repression. Because the genes required for excision and replication of the prophage are expressed only after successful removal of the CI repressor (50,58), any mutant defective in CI degradation or processing would be unable to excise and replicate following SOS induction. We first tested whether any phage replication occurred after SOS-induction in the two mutant backgrounds using Southern blotting (Fig 3A). While clear replication was visible in the wt strain background 60 min after MC induction, we did not observe replication in the Δ*clp*X mutant for either phage. By contrast, phage replication in the Δ*clp*P mutant was significantly delayed and reduced compared to the wt RN450 host strain (Fig. 3A). Complementation of the Δ*clp*X or Δ*clp*P mutations fully restored phage replication (Fig. 3B). Interestingly, we observed a faint band of phage DNA in the uninduced Δ*clp*P mutants suggesting that the phages might be replicating at low levels despite the lack of SOS induction.

**Fig. 3.**
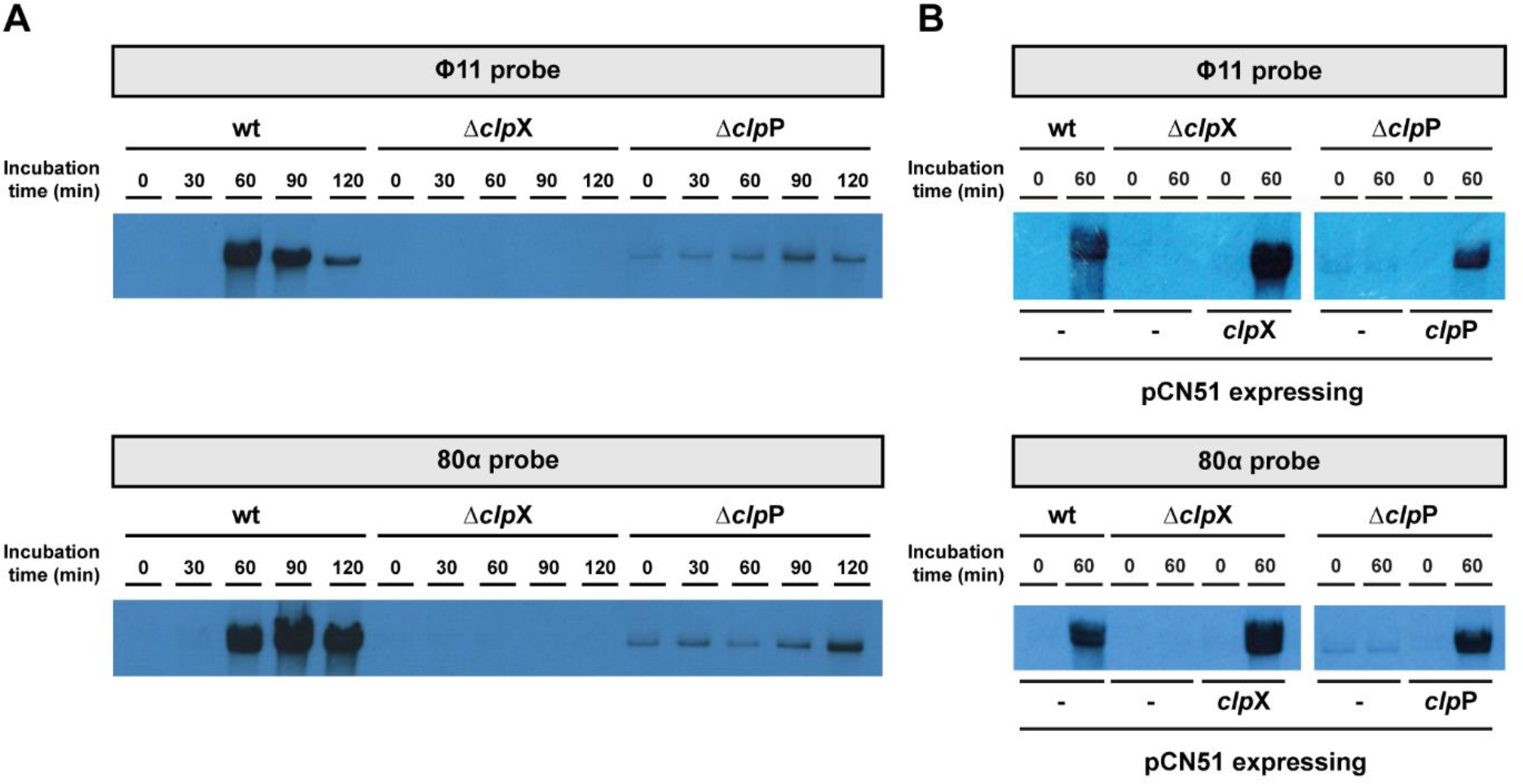
ClpP and ClpX are involved in phage induction. **(A)** The defined RN450 derivative strains lysogenic for either Φ11 (wt RN451, Δ*clp*X JP18031, Δ*clp*P JP18030) or 80α (wt JP18270, Δ*clp*X JP18169, Δ*clp*P JP18170) were induced by MC addition and samples for DNA extraction were taken at the time points indicated. Crude DNA lysates for Southern-blotting analysis were then separated by agarose gel electrophoresis, transferred onto a nitrocellulose membrane, and replicating phage DNA visualized using a phage-specific DIG-labelled DNA probe. **(B)** The *clp*X and *clp*P genes were cloned into the cadmium-inducible expression plasmid pCN51 (pJP2601 and pJP2602, respectively), introduced into the defined strains and induced by MC addition. Samples were taken at the defined timepoints for Southern blotting analysis as described in **(A)**. Expression from the pCN51 plasmids was maintained throughout the experiment by the addition of 1 μM CdCl_2_.

To have a much better picture of the roles that ClpP and ClpX play in prophage induction, we performed quantitative whole genome sequencing (50) before and 60 min after MC induction of RN450, RN450 Δ*clp*X (JP18031) or RN450 Δ*clp*P (JP18030) lysogenized with either Φ11 or 80α (Fig. 4A&B). The obtained sequencing reads were mapped to the respective strain’s genome, normalized, and graphed (Fig. 4). The wt lysogens (JP18269 for Φ11 and JP18270 for 80α), prior to induction, showed continuous sequencing coverage across both bacterial genome and prophage DNA. MC induction of these strains resulted in a sharp increase in read coverage for the prophage DNA sequence indicating replicating prophage. Note that the increased coverage of the genomic DNA to the right and left of the prophage is the result of the *in-situ* replication generated by the induced prophages (50). By contrast, and consistent with the Southern blot results (Fig. 3), phages in the Δ*clp*P mutant background (JP18158 for Φ11 and JP18170 for 80α) showed increased levels of phage DNA compared to the wt strain prior to MC induction, as indicated by the increased read coverage (Fig. 4). Interestingly, the amount of phage DNA did not increase substantially 60 min after MC induction in the Δ*clp*P mutant strain and remained considerably below replication levels of the wt strain. Importantly, neither increase nor decrease in read coverage was observed for the Δ*clp*X mutant lysogens (JP18157 for Φ11 and JP18169 for 80α) (Fig. 4). Taken together with the absence of a replicating phage band for the Δ*clp*X mutant lysogens in the Southern blots (Fig. 3A), these data therefore confirmed that ClpX was required prior to prophage excision.

**Fig. 4.**
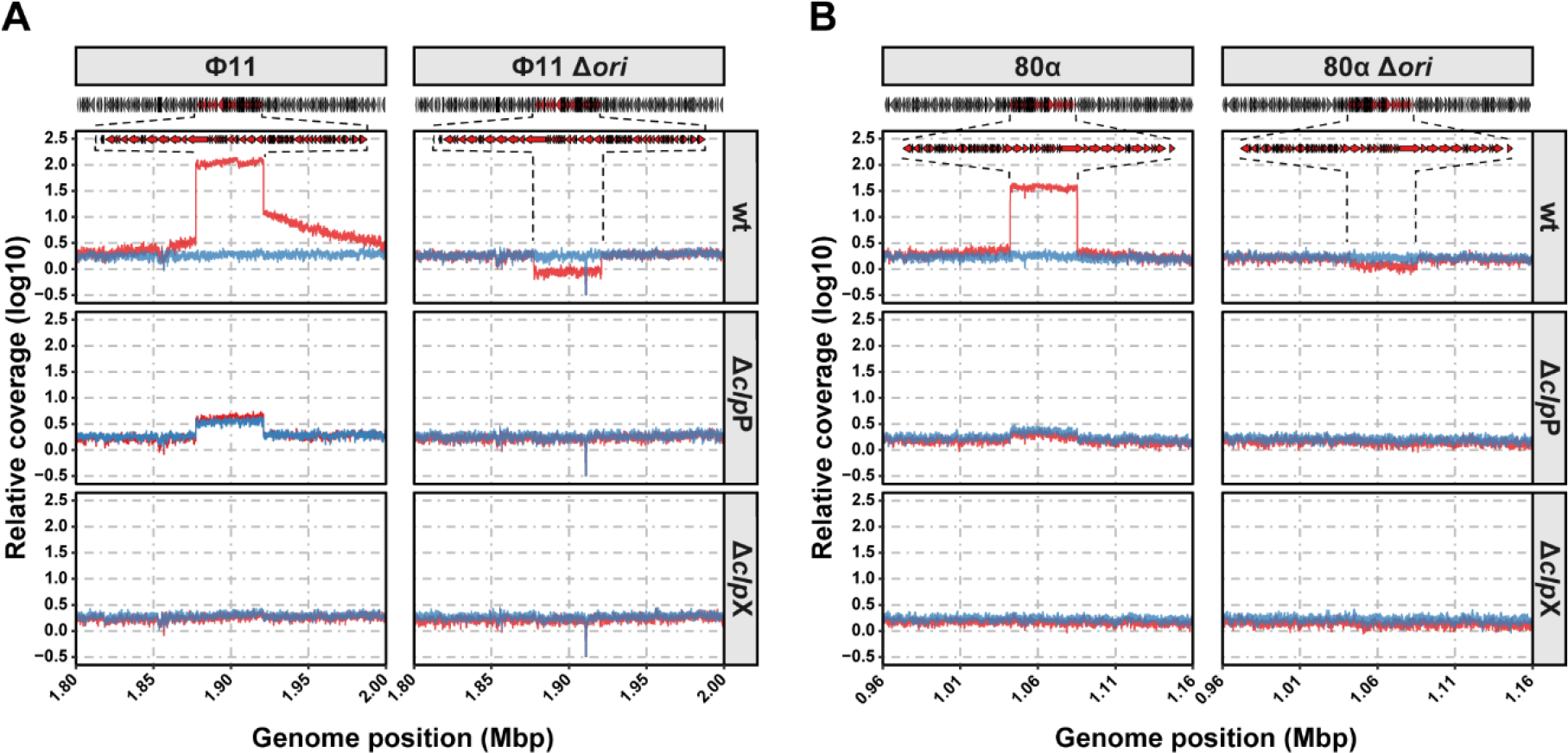
ClpP and ClpX are required for phage induction. **(A&B)** Quantitative whole genome sequencing analysis of the indicated RN450 derivative strains lysogenic for either Φ11 **(A)** or 80α **(B)** or their respective origin of replication (*ori*) mutants. The indicated strains were induced with MC and DNA isolated either before (blue) or after one hour of induction (red) with MC. DNA was then subjected to whole genome sequencing and mapped to the relevant bacterial genomes.

We also included two phage mutants in Φ11 and 80α defective their origins of replication (*ori*) as a control for a phage able to excise but not to replicate and generated Δ*clp*X and Δ*clp*P mutants in these strain backgrounds. These *ori* mutants would also pinpoint whether the Δ*clp*X or Δ*clp*P mutants were defective in phage replication or during their induction process. If ClpX and/or ClpP were involved in prophage induction (the removal and degradation of the CI repressor) or excision (excisionase is only expressed after CI degradation), these mutants would not show any changes in prophage sequence coverage before and after SOS induction in the *ori* mutant lysogens. By contrast, if either ClpX or ClpP were involved in phage replication, sequence coverage in the mutant strains in the *ori* mutant lysogens would be indistinguishable from the behavior of the *ori* mutants in the wt RN450 strain background and MC induction would lead to the loss of sequence coverage. In the wt RN450 strain background lysogenic for the Φ11 and 80α *ori* mutants (JP20045 and JP20046, respectively), the amount of prophage detected decreased substantially after MC induction, confirming that these prophage mutants excised from the bacterial chromosome and then were lost due to their inability to replicate. Deletion of the *ori* containing gene in either Φ11 or 80α in the Δ*clp*P mutant background (JP20189 and JP20192, respectively) eliminated the increased read coverage for the prophage observed with the Δ*clp*P single mutant prior to MC induction and showed no difference after MC induction (Fig.4). Since read coverage of the prophage was comparable to that of the bacterial chromosome, this confirmed that ClpP acted upstream of prophage excision. Because ClpP was essential for SOS-induction (Fig. 2), this likely prevented downstream prophage induction. However, this result does not exclude that ClpP might also be involved in the removal of CI. Similarly, deletion of the *ori* in the Δ*clp*X mutant background lysogens (JP20191 and JP20190, respectively) caused no differences in sequence coverage either with or without MC induction (Fig.4). These data therefore showed that both ClpP and ClpX were required for prophage induction and acted prior to prophage excision.

### ClpX is required for the degradation of the CI N-terminal cleavage fragment

The previous results indicated that the roles of ClpX and ClpP in SOS and prophage induction were different. While both ClpX and ClpP were shown to act upstream of phage excision in the prophage induction cascade, ClpX performed different and/or additional roles compared to ClpP. Degradation of the phage repressor is the first step in prophage induction and defects there would also prevent prophage excision. We hypothesized that ClpX and/or ClpP are likely to be involved in this process and that this was likely to occur after the initial RecA*-mediated cleavage of CI. Since it was not possibly to generate phage mutants that expressed only a post-cleavage NTD of CI, we assessed the role of ClpX and ClpP using a set of reporter plasmids reconstituting the regulatory module of Φ11 in plasmid pCN41 in which the *cro* promoter was fused to a β-lactamase reporter gene (see schematic Fig. 5). Each of these plasmids also expressed one of three versions of the CI repressor: (i) a wt CI (CI_Wt_, pJP2578), (ii) a non-degradable version thereof (CI_G131E_, pJP2590) that is insensitive to SOS-induction (57,59) and (iii) a construct expressing only the post-cleavage NTD of CI (CI_G131*_, pJP2589). This last construct mimics the N-terminal fragment of CI after SOS-induced cleavage and allowed us to study both the ability of this fragment to block phage induction as well as to assess the impact of the Δ*clp*X (JP20999) and Δ*clp*P (JP19795) deletions on its processing.

**Fig. 5.**
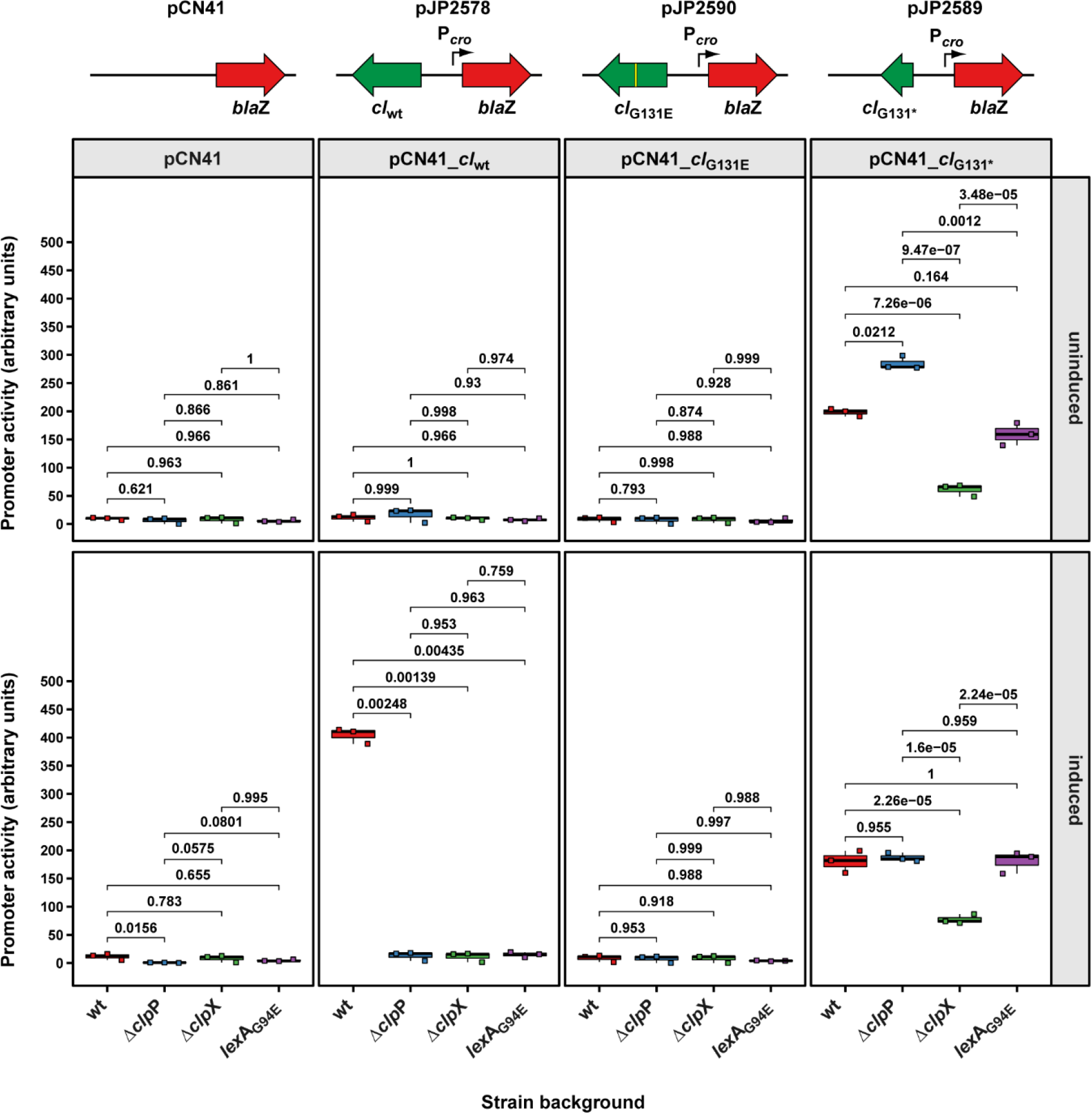
Distinct roles for ClpP and ClpX in phage induction. Plasmid pCN41-derived reporter plasmids were designed to place the β-lactamase reporter gene (*bla*Z) under the control of the Φ11 *cro* promoter. These plasmids also contained the genes encoding for either the Φ11 WT CI (*c*I_WT_), an SOS-insensitive CI mutant (*c*I_G131E_) or the post-cleavage N-terminal domain of CI alone (*c*I_G131*_). Strains containing the indicated plasmids were grown to exponential phase, split and the SOS response induced in one half of the culture with MC. Samples were taken 90 min after induction. Bold horizontal lines in each boxplot represent the median and lower and upper hinges the first and third quartiles, respectively (n=3 biological replicates). Assessment of statistically significant differences between groups was performed using ANOVA followed by Tukey’s HSD post-test. p-values are indicated above each comparison.

We introduced these reporter plasmids into the wt RN4220 strain and monitored the expression of the β-lactamase reporter either without or with MC induction. Expression of the reporter gene was repressed with the CI_Wt_ reporter plasmid (JP19910) without MC induction (Fig. 5). Addition of MC caused a significant increase in reporter gene expression levels confirming the SOS-inducibility of this construct. Reporter gene expression of the non-cleavable CI_G131E_ reporter plasmid in the wt RN4220 background (JP19912) could not be induced by MC showing its SOS-insensitivity (Fig. 5). By contrast, the reporter plasmid encoding CI_G131*_ in RN4220 (JP19911) showed high expression levels even in the absence of MC, which did not change once MC was added. This confirmed that in the wt strain background, the NTD of CI was insufficient to repress reporter expression.

We next introduced these reporter plasmids into the Δ*clp*P mutant background and repeated the experiment. The Δ*clp*P mutant carrying either the reporter plasmids expressing CI_Wt_ (JP19918) or CI_G131E_ (JP19920) showed no expression either with or without MC induction. By contrast, the CI_G131*_ reporter plasmid in this background (JP19919) highly expressed β-lactamase irrespective of the presence or absence of MC (Fig. 5). These data therefore showed that ClpP, in contrast to its role in SOS induction, was not required for the removal of the NTD fragment of CI.

When we introduced the reporter plasmids into the Δ*clp*X mutant background, we found that both the plasmid expressing CI_Wt_ (JP19914) as well as the plasmid expressing CI_G131E_ (JP19916) were fully repressed either with or without addition of MC (Fig. 5). Interestingly, the reporter plasmid expressing the NTD post-cleavage fragment CI_G131*_ (JP19915) was repressed in the Δ*clp*X mutant background and this repression could not be relieved by MC addition (Fig. 5) confirming that (i) the CI_G131*_ NTD post-cleavage fragment was still able to bind and repress the *cro* promoter and that (ii) ClpX was required for removing this repression. Both ClpX and ClpP are also involved in the initial activation of the bacterial SOS response. To determine the SOS-independent contribution of ClpX and ClpP on the processing and degradation of CI, we introduce all reporter plasmids into a strain expressing an SOS-insensitive LexA_G94E_ protein (JP1841). This mutant is no longer able to induce the bacterial SOS response and therefore is also unable to trigger the initial RecA*-dependent CI processing step in prophage activation. As expected, the CI_Wt_-expressing plasmid could no longer be induced by MC induction in this strain background (JP20406), confirming that CI was no longer cleaved by RecA*. Logically, this was also the case when introducing the SOS-insensitive CI_G131E_ reporter into the LexA_G94E_ strain background (JP20407). However, no changes compared to the wt strain were observed with the CI_G131*_ reporter plasmid (JP20408) (Fig. 5) confirming that degradation of the CI_G131*_ fragment occurred independently to any upstream SOS induction-related processes. These data therefore strongly indicated that only ClpX was essential for the processing of the CI NTD after SOS induction, while ClpP likely acted upstream through its role in SOS induction.

### The CI-NTD blocks phage infection only in the absence of ClpX

To further corroborate these observations, we circumnavigated the requirement for initial SOS induction taking advantage of the innate resistance of phage lysogens to superinfection by another phage of the same repressor type. Expression of the CI repressor from the prophage can effectively block the replication of incoming phages in a process called superinfection exclusion (Sie) (60). We reasoned that the NTD of CI would not be sufficient to block phage superinfection even when overexpressed in the wt strain background as presence of ClpX would result in its elimination. By contrast, absence of any protein required for processing the CI NTD would permit accumulation of the repressor fragment and result in a substantial reduction in phage titers after infection. To test this, we cloned the different CI alleles of Φ11 (*c*I_wt_, *c*I_G131E_ and *c*I_G131*_) into the cadmium-inducible expression vector pCN51 (pJP2584, pJP2585 and pJP2586, respectively). The plasmids were introduced into either the RN450 wt, its Δ*clp*X or Δ*clp*P mutant derivatives and the ability of Φ11 to infect these strains was assessed (Fig. 6). In the presence of pCN51 alone, the wt (JP19532), Δ*clp*P (JP19540) or Δ*clp*X (JP19536) mutant strains were susceptible to infection with Φ11 (Fig. 6). By contrast, the wt RN450 strain expressing either CI_WT_ (JP19529) or the non-degradable CI_G131E_ (JP19530) almost completely blocked phage infection, while the wt strain overexpressing the CI_G131*_ NTD (JP19531) remained susceptible to phage superinfection. Overexpression of the different CI alleles in the Δ*clp*P mutant background resulted in similar phage susceptibility levels as were observe in the wt strain background. Both CI_Wt_ (JP19537) and CI_G131E_ (JP19538) completely block Φ11 superinfection, while CI_G131*_ overexpression (JP19539) resulted in only a very marginal reduction in phage titres (Fig. 6). Similarly, both CI_Wt_ (JP19533) and CI_G131E_ (JP19534) were able to block superinfection in the Δ*clp*X mutant background. Interestingly, and consistent with a central role for ClpX in the removal of the CI NTD, overexpression of CI_G131*_ in the Δ*clp*X mutant background (JP19535) almost completely blocked phage infection. Taken together, these data confirmed that ClpX was responsible for inactivating the ability of the CI-NTD repressor fragment to block phage infection and replication, while ClpP only played a minor role in the turnover of the CI-NTD.

**Fig. 6.**
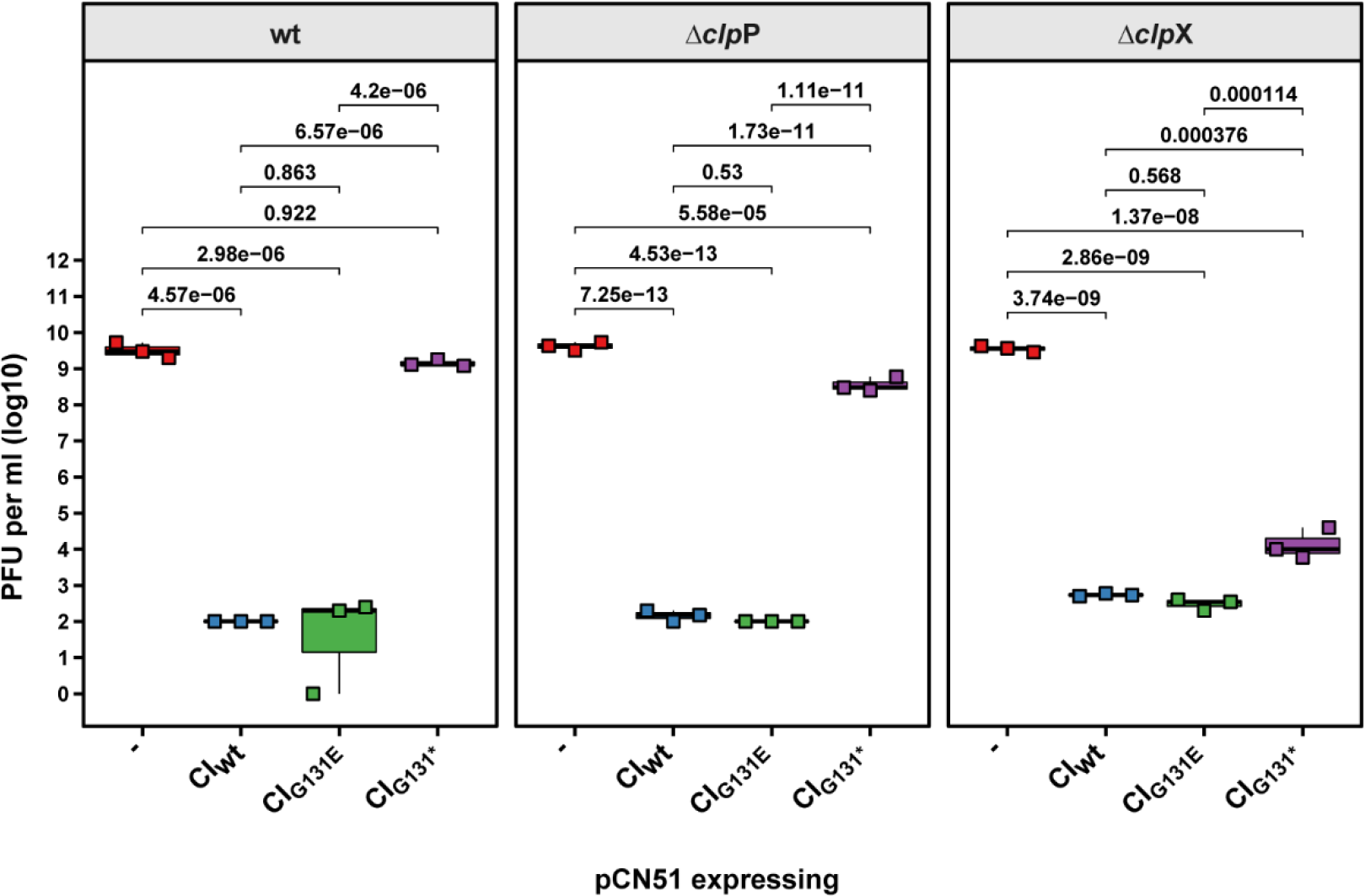
The Φ11 CI N-terminal domain affects phage infection in the absence of either ClpP or ClpX. Genes encoding either the Φ11 CI WT (CI_WT_), an SOS-insensitive version (CI_G131E_) or only its post-cleavage N-terminal fragment (CI_G131*_) were cloned into the inducible expression plasmid pCN51 and introduced into the indicated strains. Expression of from these plasmids was maintained throughout the experiment by addition of 1 μM CdCl_2_ during both growth and phage titration. Lawns of the defined, exponential phage, RN450 derivative strains were prepared on PB plates supplemented with 1 μM CdCl_2_ to maintain CI expression and serial dilutions of Φ11 lysate spotted onto these lawns. Bold horizontal lines in each boxplot represent the median and lower and upper hinges the first and third quartiles, respectively (n=3 biological replicates). Assessment of statistically significant differences between groups was performed using ANOVA followed by Tukey’s HSD post-test. p-values are indicated above each comparison.

#### ClpX-dependent prophage induction does not require interaction with ClpP

ClpX functions as a substrate specificity protein targeting individual proteins to the ClpP protease but also acts as a chaperone assisting correct protein folding. The interaction of ClpX and ClpP was previously shown to be dependent on a single amino acid and can be abolished by introducing the I265E amino acid substitution into ClpX (61). This mutant retains its chaperon function while at the same time being unable to facilitate proteolytic cleavage of protein targets. To determine whether ClpX required the subsequent interaction with ClpP for its role in phage induction, we cloned a *clp*X gene encoding either the wt or its I265E substitution (*clp*X_I265E_) into the cadmium-inducible expression vector pCN51 (pJP2601 and pJP2605, respectively) and introduced these plasmids into the Δ*clp*X mutant lysogens of Φ11 (JP18381 and JP21189, respectively) and 80α (JP18919 and JP21190, respectively). While expression of the wt ClpX protein (ClpX_wt_) in the Δ*clp*X mutant fully restored wt phage titers, expression of ClpX_I265E_ also restored phage titers to almost wt levels (Fig. 7A), indicating that proteolytic complex formation of ClpX and ClpP *per se* was not required for prophage induction. Consistent with this, expression of ClpX_I265E_ restored some, but not full, phage replication in the Δ*clp*X mutant background (Fig. 7B). Interestingly, a faint band appeared in the uninduced Δ*clp*X mutant strain expressing ClpX_I265E_ (more evident in the 80α sample) that was highly reminiscent of the band in the uninduced Δ*clp*P mutant (Fig. 3). It is possible that additional cellular processes that require ClpX and ClpP interaction are affected in this strain background, which might impact on normal phage replication. Therefore, ClpX alone is sufficient for the induction of both prophages tested and this induction does not require its interaction with ClpP.

**Fig. 7.**
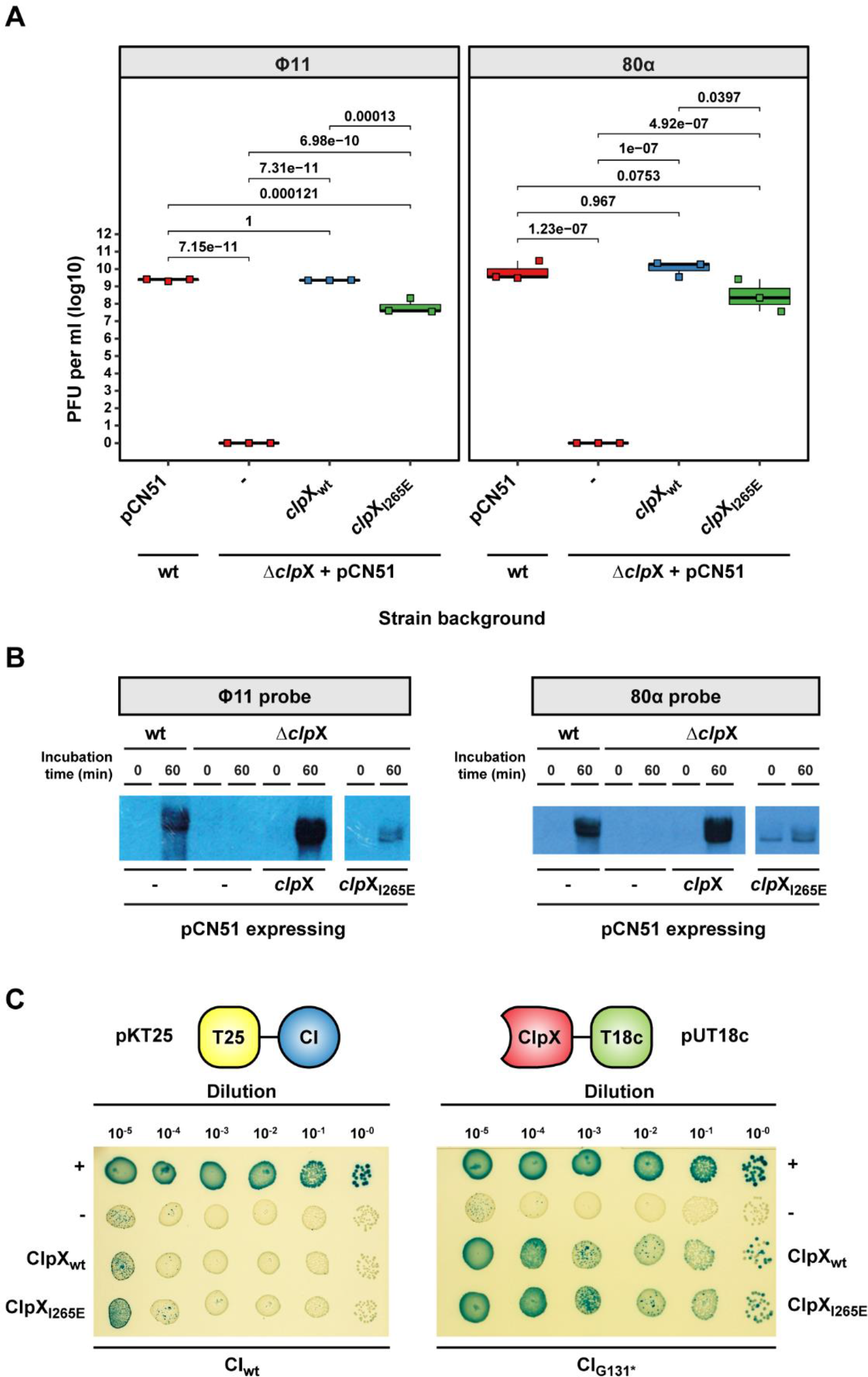
Interaction of ClpX and ClpP is not required for phage induction. **(A)** The *clp*X wt gene (*clp*X_WT_) or a *clp*X gene encoding a ClpX mutant unable to interact with ClpP (ClpX_I265E_) were cloned into the inducible expression plasmid pCN51, introduced into either the RN4220 wt or its Δ*clp*X mutant derivative lysogenic for Φ11 or 80α and induced by MC. Expression of from these plasmids was maintained throughout the experiment by addition of 1 μM CdCl_2_. Plaque formation was assessed on a lawn of RN4220. Bold horizontal lines in each boxplot represent the median and lower and upper hinges the first and third quartiles, respectively (n=3 biological replicates). Assessment of statistically significant differences between groups was performed using ANOVA followed by Tukey’s HSD post-test. **(B)** Samples of the same strains as in **(A)** were taken for DNA extraction at the time points indicated. Crude DNA lysates for Southern-blotting analysis were then separated by agarose gel electrophoresis, transferred onto a nitrocellulose membrane, and replicating phage DNA visualised using a phage-specific DIG-labelled DNA probe. **(C)** Bacterial Two-Hybrid assay of either the Φ11 full-length CI protein (CI_WT_) or the post-cleavage CI N-terminal domain only (CI_G131*_). The gene encoding either the Φ11 full-length CI protein (CI_WT_) or **(C)** the post-cleavage CI N-terminal domain only (CI_G131*_) were cloned into pKT25 (pJP2636 or pJP2632, respectively), while genes encoding either WT *clp*X (*clp*X_WT_) or ClpX unable to interact with ClpP (*clp*X_I265E_) were cloned into pUT18c (pJP2642, pJP2638, respectively). The pUT18c- and pKT25-derivative plasmids were co-transformed into *E. coli* strain BTH101 and a single colony selected. Serial dilutions of an overnight culture were plated onto LB supplemented with kanamycin (30 μg ml^−1^), ampicillin (100 μg ml^−1^), 100 μM isopropyl β-d-1-thiogalactopyranoside (IPTG) and 20 μg ml^−1^ X-gal. BTH101 transformed with pUT18c-zip and pKNT25-zip or pUT18c and pKT25 served as positive or negative controls for protein–protein interactions, respectively.

**Fig. 8.**
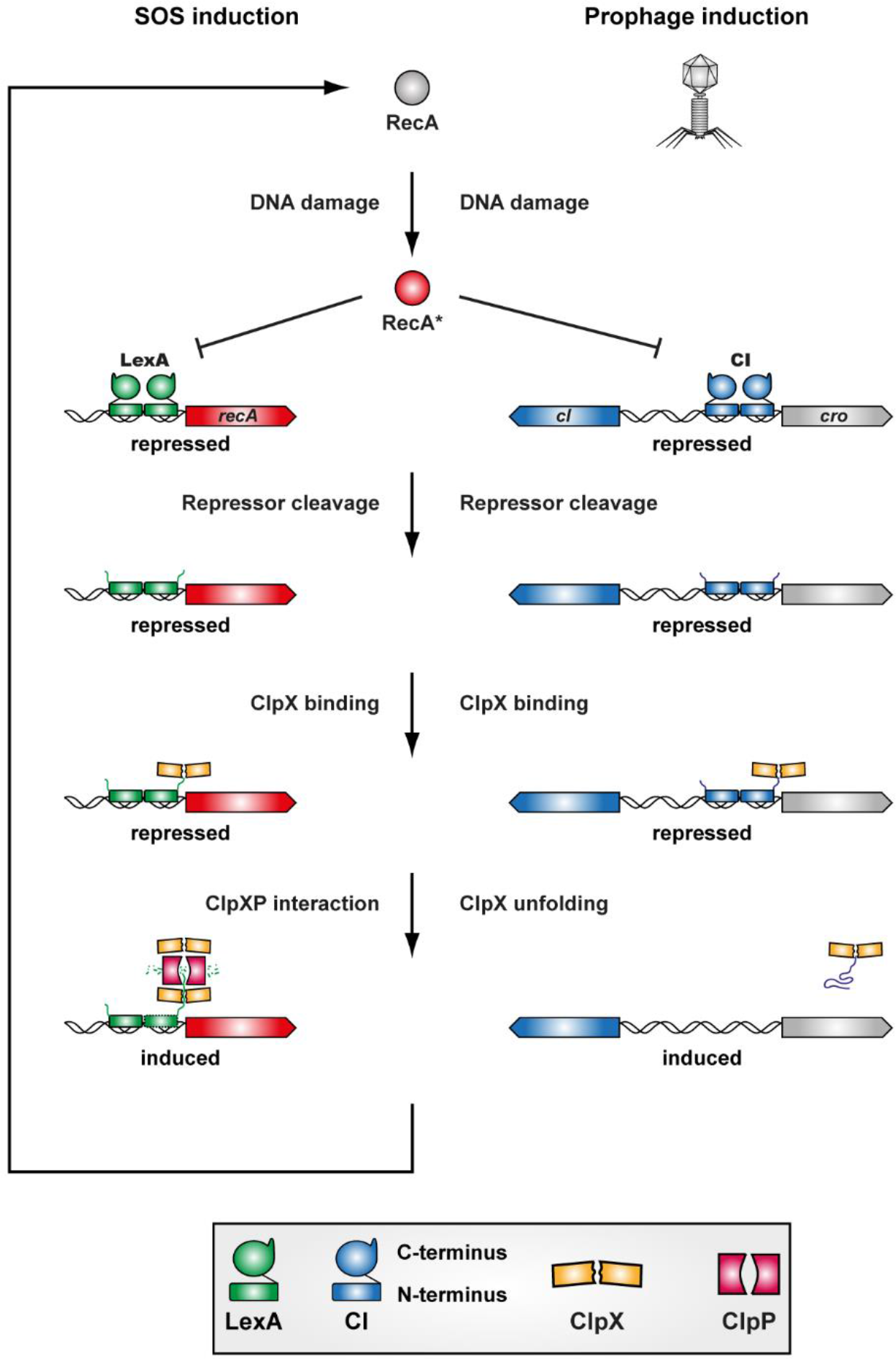
Induction of the staphylococcal SOS response and prophages. DNA damage activates the RecA (RecA*) protein which binds to both the LexA and the CI repressor catalyzing their autocleavage. The repressor N-terminal domains retain some DNA binding capacity. ClpX specifically binds to the N-terminal repressor domains after RecA* catalyzed cleavage. For SOS induction, ClpX needs to interact with ClpP to facilitate the proteolytic degradation of the LexA N-terminus. This then also results in the increased expression of RecA further increasing SOS induction for as long as DNA damage is present. By contrast, prophage induction does not require the interaction of ClpX and ClpP and binding of ClpX to the N-terminal fragment of CI is sufficient for inducting the lytic phage cycle.

#### ClpX binds to the CI N-terminal fragment but not the full-length CI repressor

Our previous results showed that the CI NTD cleavage fragment was sufficient for blocking prophage induction and infection in the absence of ClpX confirming that it retained repressor functionality. Furthermore, ClpX but not ClpP was required to relieve this repression suggesting that ClpX could interact with the N-terminal CI fragment but not the full-length protein. To investigate if this was indeed the case, we used a Bacterial Two-Hybrid system where we cloned CI_Wt_ or CI_G131*_ as C-terminal fusion products to the T25 fragment of the *Bordetella pertussis* adenylate cyclase in plasmid pKT25 (pJP2636 and pJP2632, respectively) and the ClpX_wt_ or ClpX_I265E_ mutant as N-terminal fusion product to the T18 fragment of the *B. pertussis* adenylate cyclase in plasmid pUT18c (pJP2642 and pJP2638, respectively) see schematic Fig. 7C). While ClpX did not substantially interact with CI_Wt_, interaction with its NTD alone was noticeably increased (Fig. 7C) confirming that ClpX acted after SOS-triggered CI-autocleavage. The ClpX_I265E_ mutant protein unable to interact with ClpP behaved identically to the ClpX_wt_ protein and did not interact with the CI_Wt_ but only with its separated NTD, CI_G131*_ (Fig. 7C). Taken together these data show that ClpX binds to the CI_G131*_ NTD and not the CI_Wt_ full-length protein and that this binding is sufficient to trigger phage induction even in the absence of ClpP-mediated proteolytic degradation.

## Discussion

Temperate bacteriophages can persist either in a lytic, replicative state or they can be integrated into their host’s chromosome during their lysogenic life cycle as prophages. During the lysogenic life cycle, the prophage replication genes are repressed by a master repressor, in this case CI. Activation of prophages allowing them to enter the lytic phage cycle is a fundamental process in biology that affects bacterial populations, horizontal gene transfer and evolution (62). Despite its fundamental nature, the mechanisms by which CI is fully eliminated to alleviate prophage repression has not been comprehensively resolved to date. The CI repressor shares similarity in its domain architecture and processing to the LexA repressor of the SOS response (see Fig. S3). Inactivation of both repressors is facilitated initially through their interaction with activated RecA* caused by DNA damage (17,63). This results in the autocatalytic cleavage of the repressors and separates them into a DNA-binding NTD and a CTD required for repressor dimerization. Importantly, both LexA (14–16) and CI (20–23) NTDs retain their ability to bind and repress their target promoters and require additional factors to alleviate repression. Previous studies have shown that ClpX and ClpP were both involved in the removal of the NTD of the LexA SOS-response repressor (12,14,28). The removal of the NTD of the LexA repressor is crucial for the full activation of the SOS response as it retains the ability to bind and to repress SOS-genes (14–16). Here we show, for the first time, that the ClpX ATPase activates phage replication by eliminating the CI NTD. By contrast, the ClpP proteolytic subunit primarily acts through its essential role in SOS induction rather than CI repressor degradation in prophage induction. The divergent roles for ClpP in SOS- and prophage induction might be indicative of their role in bacterial physiology and the probabilistic fate of the bacterial cell. The SOS response system is triggered to save the bacterial host and slow down physiological processes while inducing DNA repair mechanisms (6). ClpP-driven LexA-NTD turnover might be a crucial component in returning the bacterial cell to normal growth once DNA damage has been resolved. By contrast, prophage induction occurs only after the cell has undergone substantial, potentially irreparable DNA damage (63). Thus, it might be evolutionary favourable for the phage to abandon its host under such conditions. Lack of ClpP-driven degradation of the CI NTD could help to lock in the activation of the phage lytic cycle and prevent the formation of intermediary activation states that would limit the number of phage progeny released.

ClpX specifically interacts with the CI NTD after SOS-induced cleavage and not with the full-length CI proteins. Furthermore, the use of a ClpX protein unable to shuttle substrates for proteolytic degradation but able to perform its chaperone function (61) confirmed that ClpX-dependent prophage induction was independent of the proteolytic degradation of the CI N-terminal fragment. This observation is consistent with the known ability of ClpX to bind and unfold certain protein substrates such as casein in the absence of ClpP (25) as well as the role of unfoldase function of ClpX in the lifecycle of several MGEs. For example, ClpX is known to interact with the phage Mu transposome complex, where its chaperone/unfoldase activity selectively destabilizes the complex without causing its degradation (64). Similarly, the ClpX unfoldase activity can activate the plasmid replication initiation factor TrfA (65). By contrast, the phage λ O protein, a replication initiator protein which binds in complex with the λ P and the *E. coli* DnaB proteins to the phage origin of replication as a so called preprimosome, is rapidly degraded by the association of ClpX and ClpP when not in complex but is protected from proteolysis when forming part of the preprimosome (66). Degradation of free O protein is thought to influence the lysis-versus-lysogeny decision of phage λ with rapid degradation of the O-protein under certain physiological conditions favoring establishment of lysogeny rather than progression to the lytic cycle (67). Presently, only the Mu repressor protein, which is different to LexA-like CI repressors, has been shown to be subject to ClpXP mediated degradation. Even though the precise mechanism of this induction is still unknown, it is thought the a C-terminal ClpX recognition motif that is present but conformationally inaccessible in the repressed state can be exposed by environmental triggers and thus lead to the degradation of the Mu repressor and subsequent activation of the Mu transposase (68–70). The dual roles of the ClpX protein therefore necessitate the precise determination on whether the biological effects caused by ClpX are mediated via its chaperone or protein degradation function.

ClpP on the other hand was shown to be essential for the activation of the bacterial SOS response, thereby inhibiting full phage induction. Despite the inability of a *clp*P mutant to induce the SOS response, we still observed higher lysis, phage replication and higher phage titres compared to a *clp*X mutant, where phage replication was completely absent. Interestingly, loss of *clp*P gave rise to a faint phage replication band during Southern blot analysis and to increased sequence coverage during whole genome sequencing of the prophage region even in the absence of MC induction. This might suggest that ClpP performs an additional role either required for full prophage stabilization in the host’s chromosome or in preventing erroneous excision/induction events. The slow prophage induction in the *clp*P mutant might be explained by the residual expression of RecA, which can be activated by DNA damage and facilitate CI cleavage. Since RecA expression also responds to SOS induction (7) and this process is blocked in the absence of ClpP, activation of base levels of RecA could account for this slow activation.

In summary, we show that both ClpX and ClpP are involved in prophage induction in *S. aureus*. Prior to SOS-induction, CI is protected from the actions of ClpX, which only acts after SOS-mediated autocatalytic cleavage of CI by binding to the resulting NTD post-cleavage fragment and inactivating its DNA-binding capacity. While this function of ClpX alone is sufficient for prophage induction, ClpP might still be involved in degrading the CI N-terminal fragment to liberate ClpX, but this is not required for prophage induction. Instead, ClpP primarily affects prophage induction via its essential role in the staphylococcal SOS response.

## Data Availability

All underlying data are either provided within the manuscript. Reference sequences and whole genome sequencing reads can be accessed through Bioproject (https://www.ncbi.nlm.nih.gov/bioproject/) PRJNA835099.

## Acknowledgements

*Author contributions*: A.F.H. and J.R.P. conceived the study and M.A.T. conducted the experiments. M.A.T., J.R.P. and A.F.H. analyzed the data. A.F.H. wrote the initial manuscript and all authors contributed to its revisions. Funding was acquired by J.R.P.

