## Supplementary tables and figures for "The ClpX protease is essential for removing the CI master repressor and completing prophage induction in *Staphylococcus aureus*"

**Table S1 Clp proteases in *S. aureus***

| Clp protein | Locus tag (NCTC8325) | Accession number | Type |
|---|---|---|---|
| **ClpP** | SAOUHSC_00790 | YP_499347 | Protease |
| **ClpX** | SAOUHSC_01778 | YP_500283 | ATPase |
| **ClpC** | SAOUHSC_00505 | YP_499078 | ATPase |
| **ClpB** | SAOUHSC_00912 | YP_499465 | ATPase |
| **ClpL** | SAOUHSC_02862 | YP_501318 | ATPase |
| **ClpQ** | SAOUHSC_01225 | YP_499761 | Protease |
| **ClpY** | SAOUHSC_01226 | YP_499762 | ATPase |

**Table S2 Bacterial strains used in this study**

| Strain name | Relevant characteristics | Reference |
|---|---|---|
| ***Escherichia coli*** **strains** | | |
| BTH101 | Reporter strain for BACTH; F-, *gal* E15, *gal* K16, *mcr*A1, *mcr*B1, *araD139*, *rpsL1*, *hsdR2*, *cya-99* (adenylate cyclase deficient) $Str^R$) | Euromedex |
| DC10B | *mcrA* Δ(*mrr-hsdRMS-mcrBC*) φ80*lacZ*ΔM15 Δ*lacX*74 *recA*1 *araD*139 Δ(*ara-leu*)7697 *galU galK rpsL endA1 nupG* Δ*dcm* | (1) |
| DH5α | *fhuA2 lac(del)U169 phoA glnV44 Φ80' lacZ(del)M15 gyrA96 recA1 relA1 endA1 thi-1 hsdR17* | Bethesda Research Laboratories |
| IM01B | *mcrA Δ(mrr-hsdRMS-mcrBC) φ80lacZΔM15 ΔlacX74 recA1 araD139 Δ(ara-leu)7697 galU galK rpsL endA1 nupG Δdcm ΩPhelp-hsdMS (CC1-2) ΩPN25-hsdS (CC1-1)* | (2) |
| JP14175 | DH5α carrying pUT18c | This study |
| JP14176 | DH5α carrying pUT18c-zip | “ |
| JP14178 | DH5α carrying pKT25-zip | “ |
| JP14177 | DH5α carrying pKT25 | “ |
| JP15397 | DH5α carrying pCN41 | “ |
| JP16925 | DH5α carrying pJP2668 | “ |
| JP16926 | DH5α carrying pJP2669 | “ |
| JP16932 | DH5α carrying pJP2601 | “ |
| JP16933 | DH5α carrying pJP2603 | “ |
| JP16934 | DH5α carrying pJP2602 | “ |
| JP16935 | DH5α carrying pJP2604 | “ |
| JP19036 | DH5α carrying pJP2578 | “ |
| JP19042 | DH5α carrying pJP2584 | “ |
| JP19043 | DH5α carrying pJP2585 | “ |
| JP19044 | DH5α carrying pJP2586 | “ |
| JP19409 | DH5α carrying pJP2589 | “ |
| JP19410 | DH5α carrying pJP2590 | “ |
| JP20330 | DH5α carrying pJP2597 | “ |
| JP20829 | DH5α carrying pJP2596 | “ |
| JP20979 | DC10B carrying pJP2605 | “ |
| JP21575 | IM01B carrying pJP2638 | “ |
| JP21576 | IM01B carrying pJP2642 | “ |
| JP21614 | IM01B carrying pJP2636 | “ |
| JP21635 | IM01B carrying pJP2632 | “ |
| JP22129 | BHT101 carrying pUT18c-zip and pKT25-zip | “ |
| JP22130 | BHT101 carrying pUT18c and pKT25 | “ |
| JP22131 | BHT101 carrying pJP2632 and pJP2642 | “ |
| JP22132 | BHT101 carrying pJP2632 and pJP2638 | “ |
| JP22134 | BHT101 carrying pJP2642 and pJP2636 | “ |

| Strain name | Relevant characteristics | Reference |
|---|---|---|
| JP22135 | BHT101 carrying pJP2638 and pJP2636 | “ |
| ***Staphylococcus aureus strains*** | | |
| JP1361 | RN451, RN450 lysogenic for Φ11 | (3) |
| JP1841 | RN4220 expressing LexA_G94E | (4) |
| JP7834 | RN450 Δ*clp*C (internal, partial deletion of *clp*C) | (5) |
| JP7835 | RN450 Δ*clp*P (deletion of *clp*P and upstream promoter region) | (6) |
| JP7836 | RN450 Δ*clp*X (internal, partial deletion of *clp*X) | “ |
| JP7837 | RN450 *clp*L::*erm*R (plasmid disruption of *clp*L) | (5) |
| JP7838 | RN450 *clp*B::*erm*R (plasmid disruption of *clp*B) | “ |
| JP7839 | RN450 Δ*clp*QY (deletion of both genes) | (7) |
| JP8008 | JP7835 lysogenic for 80α | This study |
| JP8009 | JP7836 lysogenic for 80α | “ |
| JP8010 | JP7834 lysogenic for 80α | “ |
| JP8011 | JP7838 lysogenic for 80α | “ |
| JP8012 | JP7837 lysogenic for 80α | “ |
| JP8013 | JP7839 lysogenic for 80α | “ |
| JP8106 | JP7835 lysogenic for Φ11 | “ |
| JP8107 | JP7836 lysogenic for Φ11 | “ |
| JP8108 | JP7834 lysogenic for Φ11 | “ |
| JP8109 | JP7838 lysogenic for Φ11 | “ |
| JP8110 | JP7837 lysogenic for Φ11 | “ |
| JP8111 | JP7839 lysogenic for Φ11 | “ |
| JP18030 | RN450 Δ*clp*P | “ |
| JP18031 | RN450 Δ*clp*X | “ |
| JP18157 | RN450 Δ*clp*X, Lysogen with Φ11 | “ |
| JP18158 | RN450 Δ*clp*P, Lysogen with Φ11 | “ |
| JP18169 | RN450 Δ*clp*X, Lysogen with 80α | “ |
| JP18170 | RN450 Δ*clp*P, Lysogen with 80α | “ |
| JP18269 | RN450, Lysogen with Φ11 | “ |
| JP18270 | RN450, Lysogen with 80α | “ |
| JP18381 | JP18157 carrying pJP2601 | “ |
| JP18382 | JP18157 carrying pJP2603 | “ |
| JP18383 | JP18158 carrying pJP2602 | “ |
| JP18384 | JP18158 carrying pJP2604 | “ |
| JP18385 | JP18157 carrying pCN51 | “ |
| JP18386 | JP18158 carrying pCN51 | “ |
| JP18670 | JP18269 carrying pCN51 | “ |
| JP18916 | JP18270 carrying pCN51 | “ |
| JP18917 | JP18169 carrying pCN51 | “ |

| Strain name | Relevant characteristics | Reference |
|---|---|---|
| JP18918 | JP18170 carrying pCN51 | “ |
| JP18919 | JP18169 carrying pJP2601 | “ |
| JP18920 | JP18169 carrying pJP2603 | “ |
| JP18921 | JP18170 carrying pJP2602 | “ |
| JP18922 | JP18170 carrying pJP2604 | “ |
| JP19529 | RN450 carrying pJP2584 | “ |
| JP19530 | RN450 carrying pJP2585 | “ |
| JP19531 | RN450 carrying pJP2586 | “ |
| JP19532 | RN450 carrying pCN51 | “ |
| JP19533 | JP18031 carrying pJP2584 | “ |
| JP19534 | JP18031 carrying pJP2585 | “ |
| JP19535 | JP18031 carrying pJP2586 | “ |
| JP19536 | JP18031 carrying pCN51 | “ |
| JP19537 | JP18030 carrying pJP2584 | “ |
| JP19538 | JP18030 carrying pJP2585 | “ |
| JP19539 | JP18030 carrying pJP2586 | “ |
| JP19540 | JP18030 carrying pCN51 | “ |
| JP19795 | RN4220 Δ*clp*P | “ |
| JP19910 | RN4220 carrying pJP2578 | “ |
| JP19911 | RN4220 carrying pJP2589 | “ |
| JP19912 | RN4220 carrying pJP2590 | “ |
| JP19913 | RN4220 carrying pCN41 | “ |
| JP19914 | JP20999 carrying pJP2578 | “ |
| JP19915 | JP20999 carrying pJP2589 | “ |
| JP19916 | JP20999 carrying pJP2590 | “ |
| JP19917 | JP20999 carrying pCN41 | “ |
| JP19918 | JP19795 carrying pJP2578 | “ |
| JP19919 | JP19795 carrying pJP2589 | “ |
| JP19920 | JP19795 carrying pJP2590 | “ |
| JP19921 | JP19795 carrying pCN41 | “ |
| JP20045 | RN450 lysogen with Φ11 ΔORF15/*ori* | “ |
| JP20046 | RN450 lysogen with 80α ΔORF20/*ori* | “ |
| JP20189 | RN450 Δ*clp*P lysogen with Φ11 ΔORF15/*ori* | “ |
| JP20190 | RN450 Δ*clp*X lysogen with 80α ΔORF20/*ori* | “ |
| JP20191 | RN450 Δ*clp*X lysogen with Φ11 ΔORF 15/*ori* | “ |
| JP20192 | RN450 Δ*clp*P lysogen with 80α ΔORF20/*ori* | “ |
| JP20403 | JP1841 carrying pCN41 | “ |
| JP20406 | JP1841 carrying pJP2578 | “ |
| JP20407 | JP1841 carrying pJP2590 | “ |
| JP20408 | JP1841 carrying pJP2589 | “ |

| Strain name | Relevant characteristics | Reference |
|---|---|---|
| JP20416 | JP18031 carrying pJP2605 | “ |
| JP20417 | RN4220 carrying pJP2597 | “ |
| JP20418 | JP1841 carrying pJP2597 | “ |
| JP20419 | JP19795 carrying pJP2597 | “ |
| JP20420 | JP20999 carrying pJP2597 | “ |
| JP20858 | RN4220 carrying pJP2596 | “ |
| JP20859 | JP1841 carrying pJP2596 | “ |
| JP20860 | JP19795 carrying pJP2596 | “ |
| JP20861 | JP20999 carrying pJP2596 | “ |
| JP20999 | RN4220 Δ*clp*X | “ |
| JP21189 | JP18157 carrying pJP2605 | “ |
| JP21189 | JP18157 carrying pJP2605 | “ |
| JP21190 | JP18169 carrying pJP2605 | “ |
| JP21190 | JP18169 carrying pJP2605 | “ |
| RN450 | NCTC8325 cured of Φ11, Φ12 and Φ13 | (3) |
| RN4220 | Restriction-defective derivate of RN450 | (8) |
| RN10359 | RN450 lysogenic for 80α | (9) |

**Table S3 Plasmids used in this study**

| Plasmids name | Relevant characteristics | Reference |
|---|---|---|
| pCN41 | β-lactamase reporter plasmid for *S. aureus*, $Erm^R$ | (10) |
| pCN51 | Cadmium-inducible expression plasmid for *S. aureus*, $Erm^R$ | “ |
| pMAD | Vector for efficient allelic replacement | (11) |
| pKT25 | Bacterial Adenylate Cyclase Two-hybrid System Kit, $Km^R$ | Euromedex |
| pUT18c | Bacterial Adenylate Cyclase Two-hybrid System Kit, $Amp^R$ | Euromedex |
| pKT25-zip | Bacterial Adenylate Cyclase Two-hybrid System Kit positive control, $Km^R$ | Euromedex |
| pUT18c-zip | Bacterial Adenylate Cyclase Two-hybrid System Kit positive control, $Amp^R$ | Euromedex |
| pJP2578 | pCN41 containing $cI_{WT}$ of Φ11 cloned between SalI/BamHI | “ |
| pJP2584 | pCN51 containing $cI_{WT}$ of Φ11 cloned between SalI/BamHI | “ |
| pJP2585 | pCN51 containing $cI_{G131E}$ of Φ11cloned between SalI/BamHI | “ |
| pJP2586 | pCN51 containing $cI_{G131*}$ of Φ11 cloned between SalI/BamHI | “ |
| pJP2589 | pCN41 containing $cI_{G131*}$ of Φ11 cloned between SalI/BamHI | “ |
| pJP2590 | pCN41 containing $cI_{G131E}$ of Φ11 cloned between SalI/BamHI | “ |
| pJP2596 | pCN41 containing *lex*A promoter cloned between SalI/BamHI | “ |
| pJP2597 | pCN41 containing *rec*A promoter cloned between SalI/BamHI | “ |
| pJP2601 | pCN51 containing *clp*X of *S. aureus* cloned between BamHI/KpnI | “ |
| pJP2602 | pCN51 containing *clp*P of *S. aureus* cloned between BamHI/KpnI | “ |
| pJP2603 | pCN51 containing *clp*X of *E. faecalis* cloned between BamHI/KpnI | “ |
| pJP2604 | pCN51 containing *clp*P of *E. faecalis* cloned between BamHI/KpnI | “ |
| pJP2605 | pCN51 containing $clpX_{I265E}$ of *S. aureus* cloned between SalI/BamHI | “ |
| pJP2632 | pKNT25 containing $cI_{G131*}$ of Φ11 cloned between BamHI/KpnI | “ |
| pJP2636 | pKNT25 containing $cI_{WT}$ of Φ11 cloned between BamHI/KpnI | “ |
| pJP2638 | pUT18c containing $clpX_{I265E}$ cloned between BamHI/KpnI | “ |
| pJP2642 | pUT18c containing *clp*X cloned between BamHI/KpnI | “ |
| pJP2668 | pMAD for the deletion of *clp*X | “ |
| pJP2669 | pMAD for the deletion of *clp*P | “ |

**Table S4 Oligonucleotides used in this study**

| Plasmid name | Oligo name | Sequence (5'-3') |
|---|---|---|
| **Sequencing** | | |
| pMAD | pMAD_F | GTCCCAATATAATCATTTATCAACTCTTTTAC |
| | pMAD_R | GAAGAATCATAATGGGGAAGGCC |
| pCN51 | pCN51_F | GGTGGTCAACTTTAGAAAAGAAGG |
| | pCN51_R | GATATCAAAATTATACATGTCAACGATAATAC |
| pCN41 | pCN41_F | GGATAACCGTATTACCGCCTTTG |
| | pCN41_R | CTCTTTGGCATGTGAACTGTTTG |
| pKT25 | pKT25-F | GCAGTTCGGTGACCAGC |
| | pKT25-R | ATGTGCTGCAAGGCGATTAAG |
| pUT18c | pUT18c-F | GATGTACTGGAAACGGTGCC |
| | pUT18c-R | CTTAACTATGCGGCATCAGAGC |
| **Mutant verification** | | |
| Δ*clp*P | ClpP-9m | GGAGAAATGGTATCAACTGG |
| | ClpP-10c | TCTGGTCAACAATGGTATTC |
| Δ*clp*X | ClpX-8c | CAATTGTATCGTCTGGGTCC |
| | ClpX-7m | GAATTAGCCGGTAAAGAAGC |
| **Southern blotting probes** | | |
| Φ11 probe | Probe 11_F | ATGCAAGACCAATCATTAAAATTAGTAAAAC |
| | Probe 11_R | GATAAGCGTGGTTATATTAAGAAGTGAATGTTAC |
| 80α probe | Probe 80a_F | GTAACAGTATCAAACACTTAAGAAAAAATTC |
| | Probe 80a_R | CATAGTGACCTCCTACCATCTCATG |
| **Plasmid construction** | | |
| pJP2668 | ClpX-16cS | ACGCGTCGACAATTTTGTCTTCTTTAGTGC |
| | ClpX-5m | CTCTTTCTGCGGAAAAGACGAACTATACGACGCAGAAGG |
| | ClpX-4c | GTCTTTTCCGCAGAAAGAGC |
| | ClpX-13mB | CGCGGATCCAAATTTAAAGAAGTCCCAGA |
| pJP2669 | ClpP-8cB | ATATCGGATCCAACTGCACCTATACCTGAACG |
| | ClpP-7m | CGTGCATATGATATATACTCAGCTTAATTG |
| | ClpP-6c | TGAGTATATATCATATGCACG |
| | ClpP-5mS | GGTACCCGGGAGCTGGAAAGTTTAATGAAGG |
| pJP2638 | ClpX_152_F_B | CTAGAGGATCCAATGTTTAAATTCAATGAAGATGAAGAAAATTTG |
| | ClpX_154_R_K | AGCTCGGTACCGAGCTTTTCACTTTTATAACACATCAATGA |
| pJP2642 | ClpX_152_F_B | CTAGAGGATCCAATGTTTAAATTCAATGAAGATGAAGAAAATTTG |
| | ClpX_156_R_K | AGCTCGGTACCGAGCTTTTCACTTTTATAACACATCAATGA |

| Plasmid name | Oligo name | Sequence (5'-3') |
| --- | --- | --- |
| pJP2636 | C1_148_F_B | CTAGAGGATCCAATGGATAAAAAAGAATTAGCGAAATTTATAG |
| | C1_151_R_K | AGCTCGGTACCTGCAATACAACTTTGCCCATTACTTTAATATT |
| pJP2632 | NTD_144_F_B | CTAGAGGATCCAATGGATAAAAAAGAATTAGCGAAATTTATAG |
| | NTD_147_R_K | CGAGCTCGGTACCTGAGCACCAGTTGCACCAC |
| pJP2578 | WT CI_F_46_S | TGCAGGTCGACTCACAATACAACTTTGCCCATTACTTTAATATTAC |
| | Delta Cro_54_R_B | CCCGGGGATCCTTCTCAACTTTATTAAATTCCATTGCATG |
| pJP2589 | MT_Delta Cro_54_R_B | CCCGGGGATCCTTCTCAACTTTATTAAATTCCATTGCATG |
| | MT_G131STOP_53_F | CTGCAGGTCGACTTAAGCACCAGTTGCACCAC |
| pJP2590 | MT_Delta Cro_54_R_B | CCCGGGGATCCTTCTCAACTTTATTAAATTCCATTGCATG |
| | WT CI_F_46_S | TGCAGGTCGACTCACAATACAACTTTGCCCATTACTTTAATATTAC |
| pJP2596 | LexA IG_F_S | CCTGCAGGTCGACGTTAATTCTCTCA TATATAGGCACTCCC |
| | LexA IG_R_B | CCGGGGATCCATTTATTGTAAAACAT CATTTTCACTCCTAGAAC |
| pJP2597 | RecA IG_F_S | GCAGGTCGACGCTTAGAACAACAAA TTAATTGTATTATCGATAAAAAT |
| | RecA IG_R_B_102 | ACCCGGGGATCCGTTGATGTAGTTGA AACTCGGC |
| pJP2584 | Phi11 C1_63_F S | TGCAGGTCGACGTAAATTTAAGGAGGTAAGAAAATGGATAAAAAAGAATTAG |
| | Phi11 C1_64_R_B | CCCGGGGATCCTCACAATACAACTTTGCCCATTACTTTAATATTAC |
| pJP2586 | Phi11 C1_63_F S | TGCAGGTCGACGTAAATTTAAGGAGGTAAGAAAATGGATAAAAAAGAATTAG |
| | Phi11 G131*_65 RB | CCCGGGGATCCTCAAGCACCAGTTGCACCAC |
| pJP2585 | Phi11 C1_63_F S | TGCAGGTCGACGTAAATTTAAGGAG GTAAGAAAATGGATAAAAAAGAATT AG |
| | Phi11 C1_64_R_B | CCCGGGGATCCTCACAATACAACTT TGCCCATTACTTTAATATTAC |
| pJP2601 | ClpX_85_F_S | CTGCAGGTCGACCAATCTAGTATAGTCTTTAACGAATAGGGG |
| | ClpX_86_R_B | ACCCGGGGATCCGAGCTTTTCACTTTTATAACACATCAATG |
| pJP2603 | ClpX_106_F_B | TCTAGAGGATCCATGTACGACAATACGGATAATAACG |
| | ClpX_107_R_K | TTACCGGTACCTTATACAGCTTCTTTATCTTTTTTGTTATAAA |
| pJP2602 | ClpP_110_F_B | CTAGAGGATCCTATGTAAAATAATGAGTAACAGTTATTACAAGGAGG |
| | ClpP_111_R_K | TTATTTTGTTTCAGGTACCATCACTTC |
| pJP2604 | ClpP_108_F_B | CTAGAGGATCCTAGGCATTCAAAGTGCTTTGTGATAG |

| Plasmid name | Oligo name | Sequence (5'-3') |
|---|---|---|
| | ClpP_109_R_K | GCTCGGTACCTTTTTAATTTAAGGCGCTACTATTTTCCATTAC |
| pJP2605 | ClpX_85_F_S | CTGCAGGTCGACCAATCTAGTATAGTCTTTAACGAATAGGGG |
| | I265E_70_R | AGAAACCCTCAACTTTTTCACCAAGACGGCGCTTAAT |
| | I265E_72_F | GTGAAAAAGTTGAGGGTTTCTCAAGCAATGAAGCTGATAAATATG |
| | ClpX_86_R_B | ACCCGGGGATCCGAGCTTTTCACTTTTATAACACATCAATG |

**Supplementary Figures and Figure Legends**

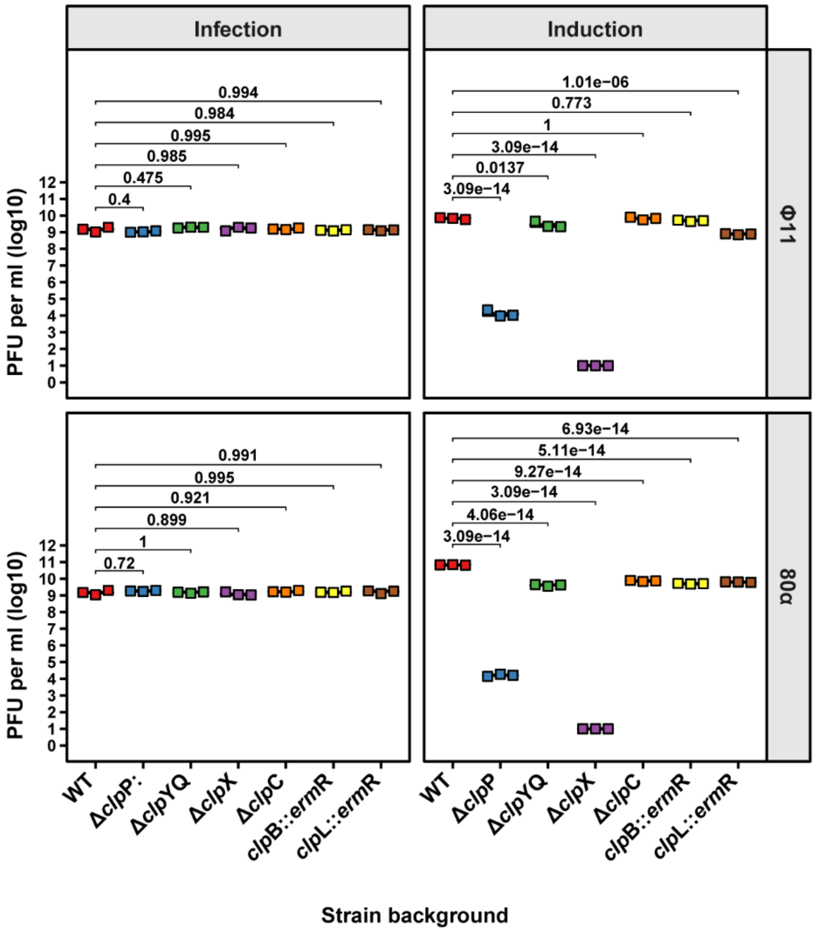


**Fig. S1. ClpP and ClpX are involved in phage induction but not phage infection.** The defined RN450 derivative mutant strains were either infected with the indicated phages or lysogenic derivatives induced by MC addition. Plaque formation was assessed on a lawn of RN4220. Bold horizontal lines in each boxplot represent the median and lower and upper hinges the first and third quartiles, respectively (n=3 biological replicates). Assessment of statistically significant differences between groups was performed using ANOVA followed by Tukey's HSD post-test. p-values are indicated above each comparison.

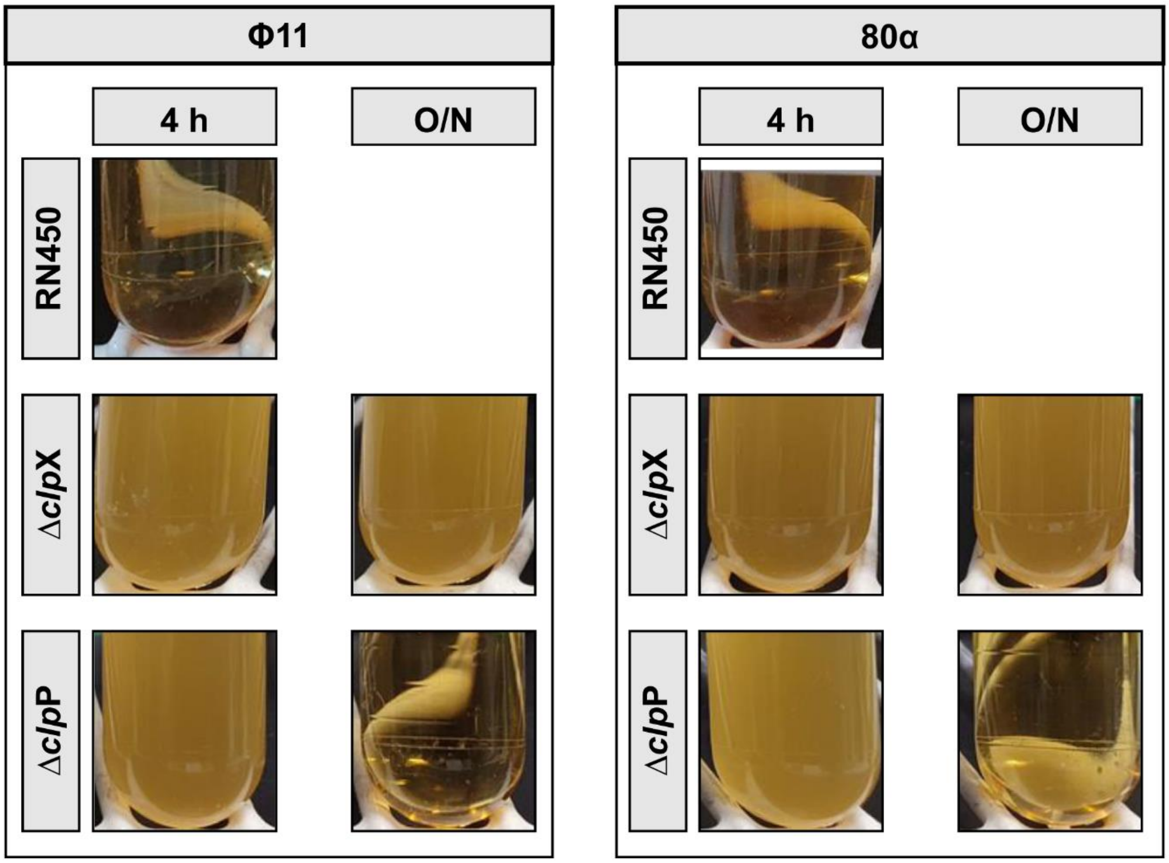


**Fig. S2. Lysis behaviour of clpX and clpP mutans.** The indicated RN450 derivatives lysogenic for either Φ11 or 80α were grown to exponential phase followed by mitomycin C induction of the lytic phage cycle. The cultures were incubated 4 h at 30℃, 80 rpm followed by an overnight incubation (O/N) on the bench. Representative images of cell lysis at different time points are shown.

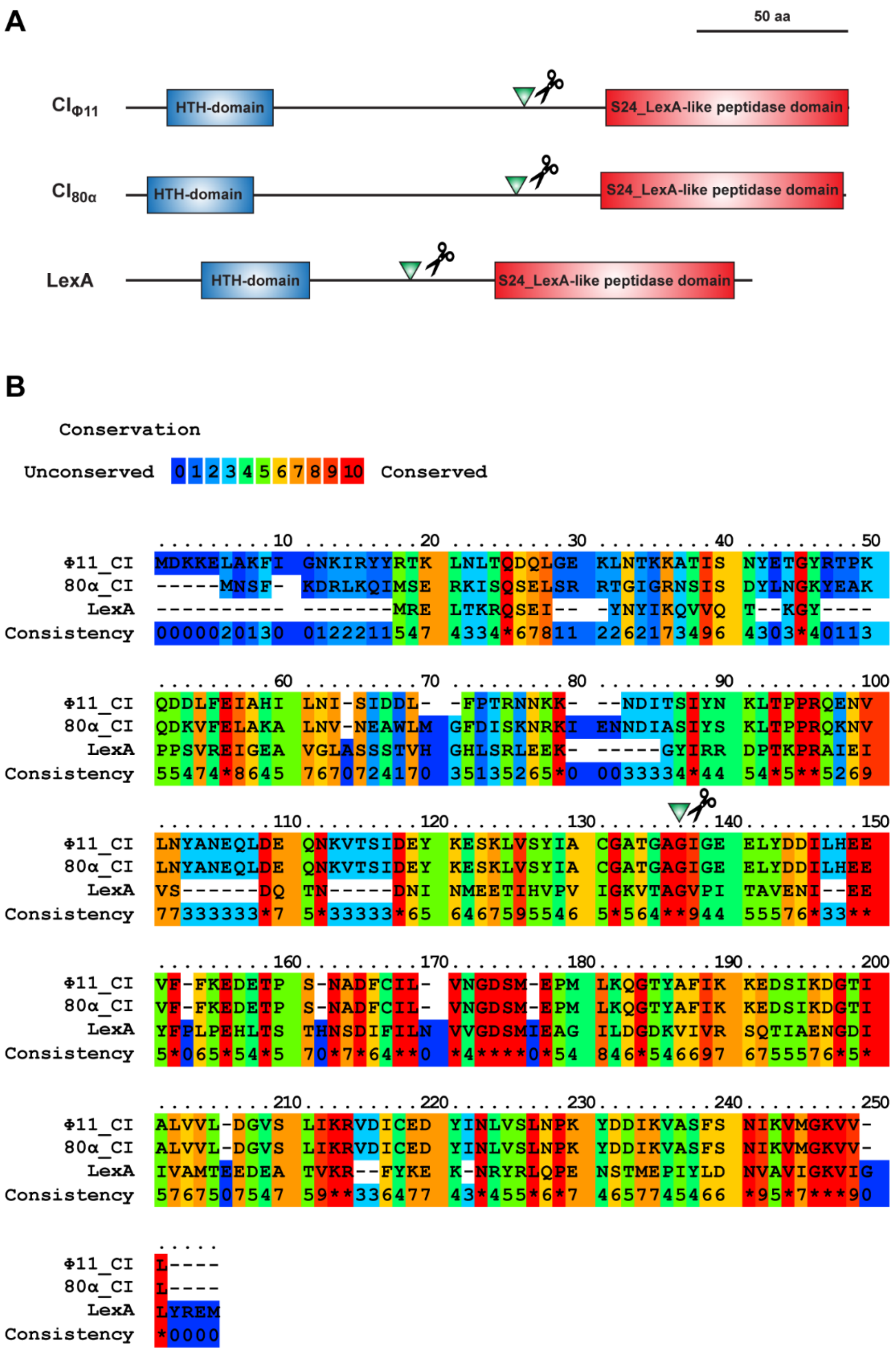


**Fig. S3. Comparison of phage repressors and LexA proteins. (A)** Schematic representation of the domain architecture of the SOS master regulator LexA with the CI repressor proteins of Φ11 and 80α. **(B)** The three proteins were aligned, and conservation of residues compared using PRALINE web server (https://www.ibi.vu.nl/programs/pralinewww/).

A green triangle accompanied by a scissors icon indicates the conserved glycine residue where cleavage occurs during RecA*-mediated autocleavage.
